# Visualizing the metazoan proliferation-terminal differentiation decision *in vivo*

**DOI:** 10.1101/2019.12.18.881888

**Authors:** Rebecca C. Adikes, Abraham Q. Kohrman, Michael A. Q. Martinez, Nicholas J. Palmisano, Jayson J. Smith, Taylor N. Medwig-Kinney, Mingwei Min, Maria D. Sallee, Ononnah B. Ahmed, Nuri Kim, Simeiyun Liu, Robert D. Morabito, Nicholas Weeks, Qinyun Zhao, Wan Zhang, Jessica L. Feldman, Michalis Barkoulas, Ariel M. Pani, Sabrina L. Spencer, Benjamin L. Martin, David Q. Matus

## Abstract

Cell proliferation and terminal differentiation are intimately coordinated during metazoan development. Here, we adapt a cyclin-dependent kinase (CDK) sensor to uncouple these cell cycle-associated events live in *C. elegans* and zebrafish. The CDK sensor consists of a fluorescently tagged CDK substrate that steadily translocates from the nucleus to the cytoplasm in response to increasing CDK activity and consequent sensor phosphorylation. We show that the CDK sensor can distinguish cycling cells in G1 from terminally differentiated cells in G0, revealing a commitment point and a cryptic stochasticity in an otherwise invariant *C. elegans* cell lineage. We also derive a predictive model of future proliferation behavior in *C. elegans* and zebrafish based on a snapshot of CDK activity in newly born cells. Thus, we introduce a live-cell imaging tool to facilitate *in vivo* studies of cell cycle control in a wide-range of developmental contexts.

## Introduction

Organismal development requires a delicate balance between cell proliferation and cell cycle arrest. In early embryos, the emphasis is placed on rapid cell proliferation, which is achieved by omitting gap phases (G1 and G2) and establishing a biphasic cell cycle that rapidly alternates between DNA synthesis (S phase) and mitosis (M phase) (Edgar and O’Farrell, 1989; Newport and Kirschner, 1982). After several rounds of embryonic cell division, the gap phases are introduced, coincident in many organisms with cell fate decisions and the execution of morphogenetic cell behaviors (Foe, 1989; Grosshans and Wieschaus, 2000). These gap phases are believed to function as commitment points for cell cycle progression decisions. The earliest point of commitment occurs during G1, which is the focus of this study. Cells either engage in cell cycle progression and enter S phase, or they exit the cell cycle altogether and enter a cell cycle arrested state referred to as G0 and undergo quiescence or terminal differentiation (Sun and Buttitta, 2017). Although the location of the G1 commitment point in yeast (Start) and cultured mammalian cells (Restriction Point) has in large part been spatiotemporally mapped and molecularly characterized (Hartwell et al., 1974; Pardee, 1974; Spencer et al., 2013), when metazoan cells make this decision *in vivo* while integrating intrinsic and the extrinsic cues of their local microenvironment during development remains poorly understood. A cell cycle sensor that is amenable to such *in vivo* studies can shed new light on this four-decade-old biological phenomenon.

In 2008, Sakaue-Sawano and colleagues engineered a multicolor fluorescent ubiquitination-based cell cycle indicator (FUCCI) for mammalian cell culture (Sakaue-Sawano et al., 2008). FUCCI has since been adapted for many research organisms (Ozpolat et al., 2017; Zielke and Edgar, 2015). However, FUCCI on its own cannot distinguish between a cell residing in G1 that will cycle again upon completing mitosis and a cell that is poised to enter G0 (Oki et al., 2014). Separating G1 from G0 is essential to understanding mechanisms controlling cell cycle exit during quiescence or terminal differentiation. To distinguish G1 from G0 in mammalian cell culture, Hanh, Spencer and colleagues developed and implemented a single-color ratiometric sensor of cell cycle state comprised of a fragment of human DNA helicase B (DHB) fused to a fluorescent protein that is phosphorylated by CDKs (Hahn et al., 2009; Schwarz et al., 2018; Spencer et al., 2013). Notably, through quantitative measurements of CDK activity, this sensor provided new insights into the proliferation-quiescence decision in cultured mammalian cells by identifying cycling cells that exit mitosis in a CDK-increasing (CDK^inc^) state and quiescent cells that exit mitosis in a CDK-low (CDK^low^) state (Spencer et al., 2013). Nonetheless, a DHB-based CDK sensor has not been utilized to evaluate the proliferation-terminal differentiation decision.

In this study, we investigate the proliferation-terminal differentiation decision in *C. elegans* and zebrafish, two powerful *in vivo* systems with radically different modes of development. We generate transgenic CDK sensor lines in each organism to examine this decision live at mitotic exit. By quantifying CDK activity, or DHB ratios, at mitotic exit, we are able to predict future cell behavior across several embryonic and post-embryonic lineages. Despite cells generally exiting mitosis with decreased CDK activity levels, we reliably distinguish cycling cells that exit mitosis into G1, in a CDK^inc^ state, from terminally differentiated cells that exit mitosis into G0, in a CDK^low^ state. To gain insights into cell cycle progression commitment, we examine the activity of *C. elegans cki-1*, a cyclin-dependent kinase inhibitor (CKI) of the Cip/Kip family, demonstrating that endogenous CKI-1 levels are anti-correlated with CDK activity during the proliferation-terminal differentiation decision. We propose that integration of CKI-1 levels in the mother cell and the high CKI-1/low CDK activity at mitotic exit mediate this decision. By utilizing the CDK sensor to predict future cell behavior, we uncover a cryptic stochasticity that occurs in a temperature-dependent fashion in the *C. elegans* vulva, an otherwise invariant and well-characterized lineage. Finally, we reveal cell cycle dynamics in zebrafish, an organism that lacks a defined cell lineage, demonstrating that terminally differentiated embryonic tissues display DHB ratios that correlate with those observed in terminally differentiated G0 cells in *C. elegans*. Together, we present a tool for visualizing G1/G0 dynamics *in vivo* during metazoan development that can be used to study the interplay between cell proliferation and terminal differentiation.

## Results

### Design and Characterization of a Live *C. elegans* CDK Sensor to Define Interphase States

We synthesized a codon-optimized fragment of human DHB comprised of amino acids 994–1087 (Hahn et al., 2009; Spencer et al., 2013). The fragment contains four serine sites that are phosphorylated by CDKs in human cells (Moser et al., 2018; Spencer et al., 2013). These serine sites flank a nuclear localization signal (NLS) situated next to a nuclear export signal (NES) (**Figure 1A**). When CDK activity is low, the NLS is dominant over the NES and DHB localizes to the nucleus. However, when CDK activity increases (i.e., during cell cycle entry), the NLS is obstructed and DHB localizes to the cytoplasm (**Figure 1B**). Using this DHB fragment, we generated two CDK sensors by fusing green fluorescent protein (GFP) or two copies of a red fluorescent protein, mKate2 (2xmKate2), to the DHB C-terminus (**Figure 1A**). To visualize the nucleus, we co-expressed *his-58*/histone H2B fused to 2xmKate2 or GFP, respectively, which is separated from DHB by a P2A self-cleaving viral peptide (Ahier and Jarriault, 2014). We drove the expression of each CDK sensor via a ubiquitous *rps-27* promoter (Ruijtenberg and van den Heuvel, 2015).

**Figure 1.**
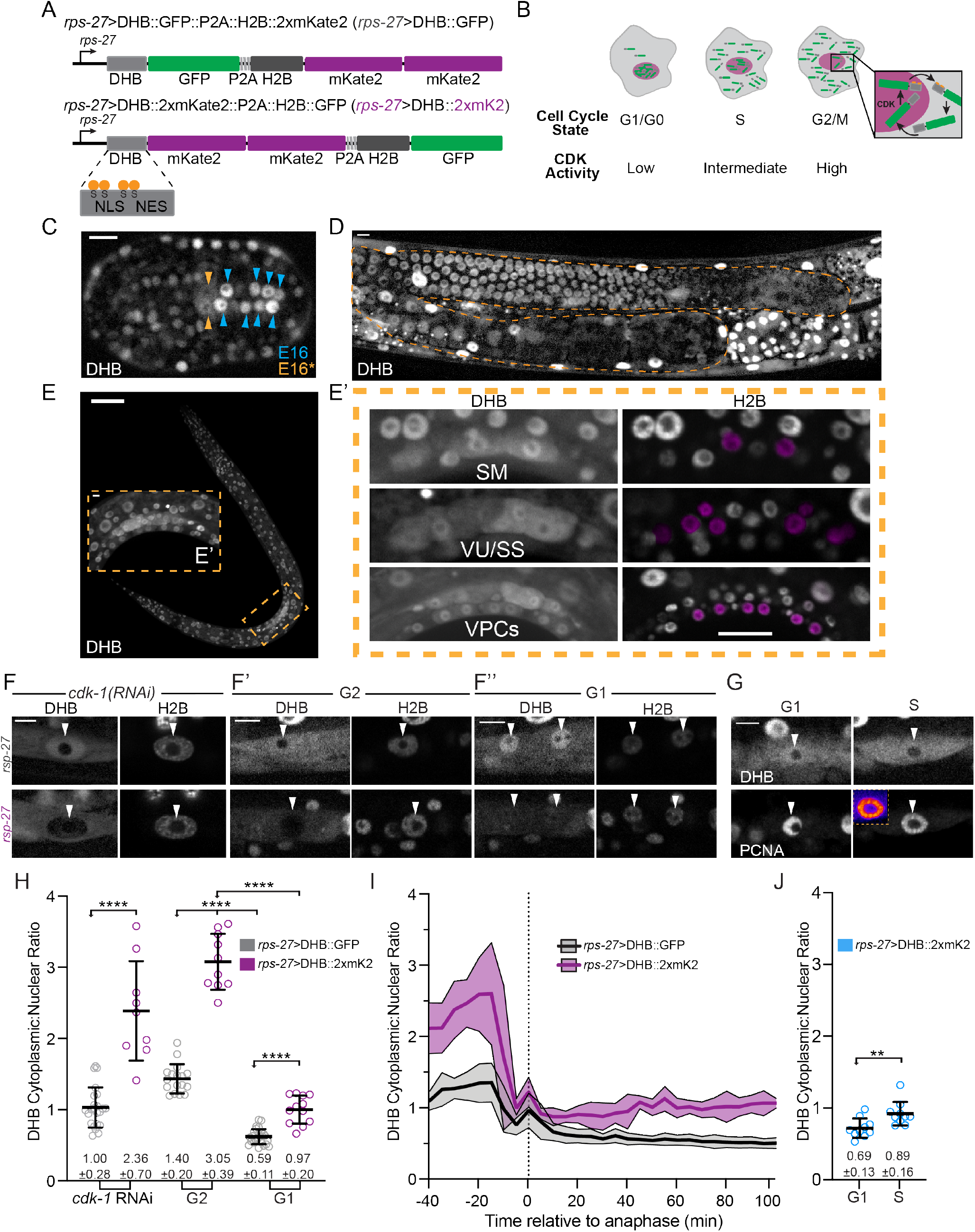
Design and characterization of a live *C. elegans* CDK sensor to define interphase states. (A) Schematic of the CDK sensor fused to GFP (top) or two copies of mKate2 (bottom) and a nuclear mask (H2B::FP) separated by a self-cleaving peptide (P2A). Inset: nuclear-localization signal (NLS), nuclear export signal (NES), and consensus CDK phosphorylation sites, serine (S) residues. (B) Schematic of CDK sensor translocation during the cell cycle. (C) Quiescent E16 cells (blue arrows) versus the cycling E16* star cells (orange arrows) arrows. (D, E) Confocal micrograph montage of CDK sensor in the *C. elegans* germline (D, orange dashed line) and L3 stage larvae (E). Three somatic tissues are highlighted (inset, dashed orange box) shown at higher magnification in E’, with pseudo-colored nuclei (magenta) depicting cells of interest. (F, G) Representative images of sensor expression in SM cells following *cdk-1* RNAi treatment, at peak G2 (F’), and 20 min after anaphase during G1 (F’’) in DHB::GFP (grey) and DHB::2xmKate2 (magenta) as well as DHB::2xmKate2 co-expressed with PCNA (*pcn-1*>PCN-1::GFP; G), inset highlights PCNA puncta in S phase. (H) Dot plot depicting dynamic ranges of the two CDK sensor variants, measured by the cytoplasmic:nuclear ratio of DHB mean fluorescent intensity, at G2 and G1 in the SMs (*n*≥9 cells). (I) Plot of DHB cytoplasmic:nuclear ratio in SM cells during one round of cell division, measured every 5 min (*n*=4 cells per strain). Dotted line indicates time of anaphase. Error bars and shaded error bands depict mean±SD. (J) Dot plot depicting range of G1 and S phase in DHB::2xmKate2 based on absence or presence of PCNA puncta (*n*=10 cells per phase). \*\**p*≤0.01, \*\*\*\**p*≤0.0001. Significance determined by statistical simulation; *p*-values in Table S1. Scale bar = 10 μm (except in E: 20 μm and F: 5 μm).

To test both the GFP (**Figure 1C, 1D, S1A**) and 2xmKate2 (**Figure S1B-D**) versions of our CDK sensor, we began by examining cell divisions in the *C. elegans* embryo and germline (**Movie S1**). First, we visualized cells in the embryonic intestine, which is clonally derived from the E blastomere, as these are the first cells in the embryo to have gap phases (Edgar and McGhee, 1988). The E blastomere goes through four rounds of divisions to give rise to 16 descendants (E16 cells) about four hours after first cleavage. While 12 of the E16 cells have completed their embryonic divisions at this stage (Leung et al., 1999), four cells called E16* star cells divide once more to generate the 20-celled intestine (E20). Although all E16 cells polarize and show signs of differentiation, the E16* star cells quickly reenter the cell cycle to divide again (Rasmussen et al., 2013; Yang and Feldman, 2015). Thus, we wondered whether our CDK sensor could be used to distinguish between cycling E16* star cells and quiescent E16 cells. To accomplish this, we tracked E16* star cell division from the E16–E20 stage and observed that DHB::GFP localizes in a cell cycle-dependent fashion during these divisions, with DHB::GFP translocating from the nucleus to the cytoplasm and then re-locating to the nucleus at the completion of E16* star cell division (**Figure 1C, S1A**). Consistent with our observations using the GFP version of our CDK sensor in mid-embryogenesis, DHB::2xmKate2 also dynamically translocates from the nucleus to the cytoplasm during cell divisions in the early embryo (**Figure S1B**). Second, we examined the localization of DHB::GFP (**Figure 1D**) and DHB::2xmKate2 (**Figure S1C, S1D**) in the adult *C. elegans* germline. Here we detected a gradient of live CDK activity, from high in the distal mitotic progenitor zone to low in the proximal meiotic regions, as described with EdU incorporation and phospho-histone H3 staining (Kocsisova et al., 2018). Together, these results demonstrate that our CDK sensor is dynamic during cell cycle progression in the *C. elegans* embryo and germline.

The ability to distinguish cycling cells from quiescent cells in the embryo made us wonder whether we could also distinguish cycling cells from terminally differentiated cells post-embryogenesis. Therefore, we examined our CDK sensor in several post-embryonic somatic lineages that undergo proliferation followed by terminal differentiation (Sulston and Horvitz, 1977). Specifically, we selected the sex myoblasts (SM), the somatic sheath (SS) and ventral uterine (VU) cells of the somatic gonad, and the vulval precursor cells (VPCs) (**Figure 1E and E’**). To define each phase of the cell cycle while these lineages are proliferating, we combined static and time-lapse imaging approaches to measure cytoplasmic:nuclear DHB ratios for G1, S and G2 (**Figure 1F-J, S1E-M**). First, we RNAi depleted the sole *C. elegans* CDK1 homolog, *cdk-1,* to induce a penetrant G2 phase arrest in the SM cells (**Figure 1F**). Quantification of DHB ratios following *cdk-1* RNAi treatment showed a mean ratio of 1.00±0.28 and 2.36±0.70 in the GFP and 2xmKate2 versions of our CDK sensor, respectively (**Figure 1H**). Next, we quantified DHB ratios following time-lapse of SM (**Figure 1F’, 1H, S1G**), uterine (**Figure S1E, S1G**) and VPC (**Figure S1F, S1G**) divisions to determine peak values of G2 CDK activity (**Figure 1I, S1H, S1I**). All lineages exhibited the same CDK sensor localization pattern during peak G2—that is, maximal nuclear exclusion. Next, for each lineage (**Figure 1F’’, S1E’, S1F’**), we quantified DHB ratios 25 min after anaphase from our time-lapses to determine a threshold for G1 phase CDK activity. In G1, DHB::GFP and DHB::2xmKate2 were nuclear localized after mitotic exit with mean ratios of 0.59±0.11 and 0.97±0.20 in SMs, 0.67±0.10 and 1.13±0.17 in uterine cells, and 0.35±0.14 and 0.58±0.32 in VPCs (**Figure 1H, 1I, S1G-I**). Finally, we paired DHB::2xmKate2 with a reporter for S phase, fusing GFP to the sole *C. elegans* proliferating cell nuclear antigen (PCNA) homolog, *pcn-1*, expressed under its own endogenous promoter at single copy. Although nuclear localized throughout the cell cycle, PCNA forms sub-nuclear puncta solely in S phase (Brauchle et al., 2003; Dwivedi et al., 2019; Strzyz et al., 2015; Zerjatke et al., 2017). Analysis of time-lapse data found that punctate expression of PCN-1::GFP correlated with mean DHB::2xmKate2 ratios of 0.89±0.16 in SM (**Figure 1G, 1J; Movie S2**), 1.00±0.10 in uterine (**Figure S1J, S1K**) and 1.02±0.22 in VPC (**Figure S1J, S1K**) lineages. Despite individual lineages showing differences in CDK activity (**Figure S1G, S1K, S1L, S1M**), primarily in G1, we can establish DHB ratios for each interphase state (G1/S/G2) across several post-embryonic somatic lineages using either CDK sensor and recommend pairing with a PCNA reporter for precise determination of interphase state boundaries. We next wondered if we could distinguish G1 from G0 as these somatic lineages exit their final cell division; therefore, allowing us to visibly and quantitatively detect terminal differentiation *in vivo*. We mainly chose the DHB::GFP version of our CDK sensor to conduct the following experiments as it was more photostable.

### CDK^low^ Activity after Mitotic Exit is Predictive of Terminal Differentiation

In asynchronously dividing MCF10A epithelial cell lines, cells that exited mitosis into a CDK2^low^ state had a high probability of staying in G0 compared to cells that exited at a CDK2^inc^ state (Spencer et al., 2013). We therefore wanted to determine whether the cytoplasmic:nuclear ratio of DHB::GFP following an *in vivo* cell division could be used to predict if a cell will enter G1 and divide again or enter G0 and terminally differentiate. Taking advantage of the predictable cell lineage pattern of *C. elegans*, we quantitatively correlated DHB::GFP ratios with the decision to proliferate or terminally differentiate. We first quantified DHB::GFP ratios from time-lapse acquisitions of SM cell divisions. The SM cells undergo three rounds of cell division during the L3 and L4 larval stages before terminally differentiating into uterine(um) and vulval muscle(vm) (**Figure 2A**) (Sulston and Horvitz, 1977). Quantification of DHB::GFP in this lineage revealed that shortly after the first and second divisions, CDK activity increases immediately after mitotic exit from an intermediate level, which we designate as a CDK^inc^ state (**Figure 2B, 2C; Movie S3**), Conversely, CDK activity following the third and terminal division remains low, designated as a CDK^low^ state. Bootstrap analyses support a significant difference in DHB::GFP ratios between pre-terminal (CDK^inc^) and terminal divisions, but not among pre-terminal divisions (**Figure S2A-C**). We then quantified DHB::GFP ratios during the division of two somatic gonad lineages, the VU and SS cells. They both undergo several rounds of division during the L3 larval stage and terminally differentiate in the early L4 stage (**Figure 2D**) (Sulston and Horvitz, 1977). We quantified a pre-terminal division and the subsequent division that leads to terminal differentiation. Similar to the SM lineage, both somatic gonad lineages exit the round of cell division prior to their final division into a CDK^inc^ state and then exit into a CDK^low^ state following their terminal differentiation (**Figure 2E, 2F, S2D-F; Movie S4**). Bootstrap analyses support a significant difference between DHB::GFP ratios in pre-terminal versus terminal divisions in the developing somatic gonad (**Figure S2D**).

**Figure 2.**
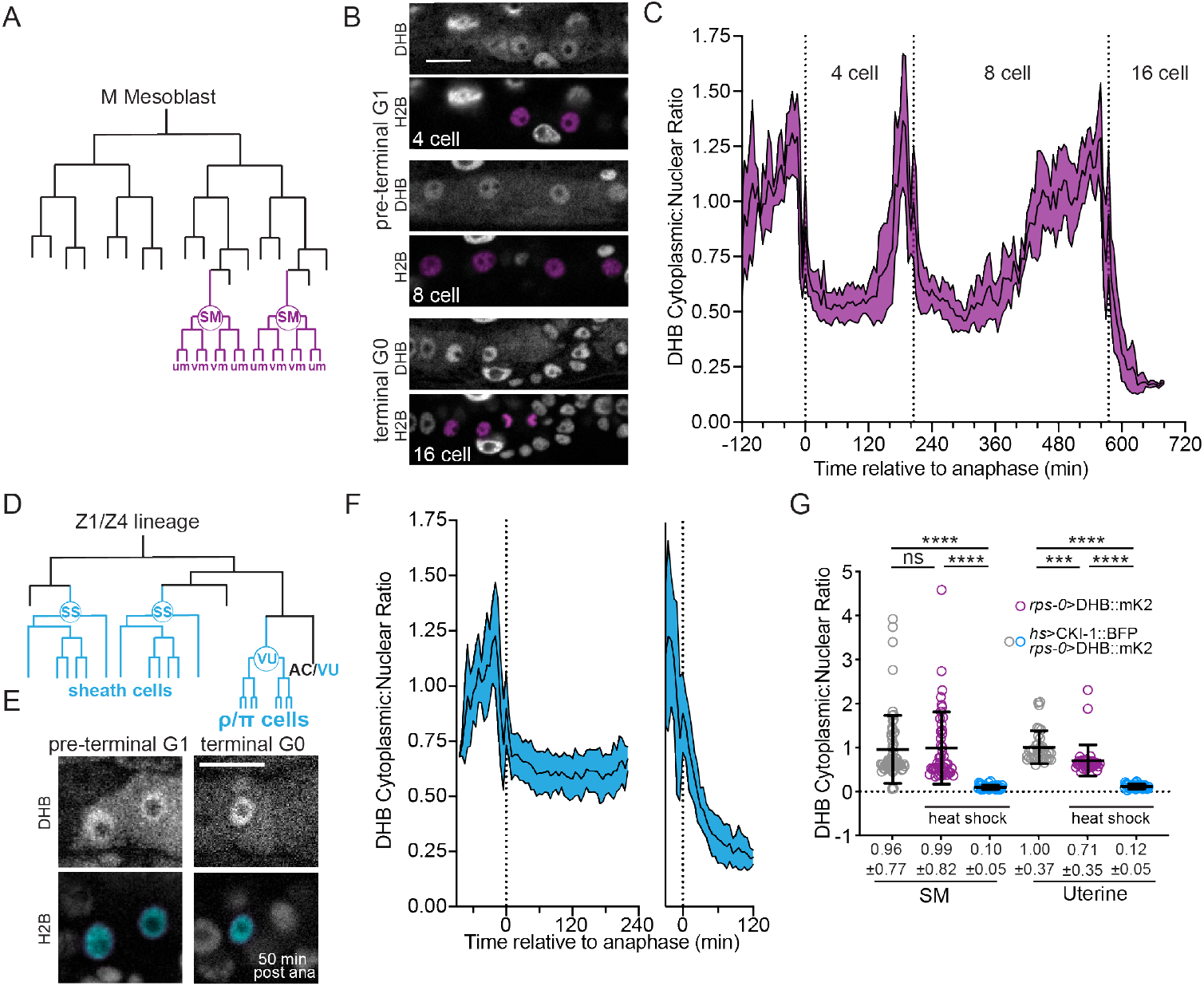
Sex myoblasts and somatic gonad cells exit terminal divisions into a CDK^low^ state. SM (A) and uterine (D) lineage schematics. (B) Micrographs of a time-lapse movie showing SM cells (B) and uterine cells (E) at G1 in pre-terminal divisions and G0 upon terminal differentiation. Quantification of CDK sensor in SM cells (C; *n*≥10) and uterine cells (F; *n*≥13). (G) Quantification of CDK sensor localization in SM cells and uterine cells following ectopic expression of CKI-1 (*hsp*>CKI-1::2xmTagBFP2) compared to non-heat shock controls and heat shock animals without the inducible transgene (*n*≥36 cells per treatment). Pseudo-colored nuclei (magenta, B; cyan, E) indicate cells of interest. Scale bars = 10 μm. Dotted line indicates time of anaphase. Line and shaded error bands depict mean±SD. Time series measured every 5 min. ns, not significant, \*\*\*\**p*≤0.0001. Significance determined by statistical simulations; *p*-values in Table S1.

Next, we sought to determine how the CDK sensor behaves under conditions in which cells are experimentally forced into G0. To accomplish this, we generated a single copy transgenic line of mTagBFP2-tagged CKI-1, the *C. elegans* homolog of p21^Cip1^/p27^Kip1^, under an inducible heat shock promoter (*hsp*), paired with a *rps-0*>DHB::mKate2 variant of the CDK sensor. Induced expression of CKI-1 is expected to result in G0 arrest (Hong et al., 1998; Matus et al., 2014; van der Horst et al., 2019). Indeed, in the SM and uterine lineages, induced expression of CKI-1 resulted in cells entering a CDK^low^ G0 state, with mean DHB ratios of 0.10±0.05 and 0.12±0.05, respectively (**Figure 2G**) as compared to control animals that lacked heat shock-induced expression (SM: 0.99±0.82, uterine: 0.71±0.35) or lacked the inducible transgene (SM: 0.96±0.77, uterine: 1.00±0.37). Thus, induced G0 arrest by ectopic expression of CKI-1 is functionally equivalent, by CDK activity levels, to the G0 arrest that occurs following mitotic exit in an unperturbed cell destined to undergo terminal differentiation.

We next examined the divisions of the 1°- and 2°-fated VPC lineage. The *C. elegans* vulva is derived from three cells (P5.p–P7.p), which undergo three rounds of cell division during the L3 and early L4 larval stages (**Figure 3A, 3B**) (Katz et al., 1995; Sternberg and Horvitz, 1986; Sulston and Horvitz, 1977). Rather than giving rise to 24 cells, the two D cells, the innermost granddaughters of the 2°-fated P5.p and P7.p, terminally differentiate one round of cell division early. This results in a total of 22 cells which comprise the adult vulva (Katz et al., 1995; Sulston and Horvitz, 1977). Quantification of DHB::GFP ratios during VPC divisions yielded the expected pattern. The daughters of P5.p–P7.p all exited their first division into a CDK^inc^ state (**Figure 3C, 3D**). After the next division, the 12 granddaughters of P5.p–P7.p (named A–F symmetrically) are born, including the terminally differentiated D cell (Katz et al., 1995; Sulston and Horvitz, 1977). At this division, the strong nuclear localization of DHB::GFP in the D cell was in stark contrast to the remaining proliferating VPCs. The D cell exited into and remained in a CDK^low^ state, while the remaining VPCs exited into a CDK^inc^ state and continued to progress through the cell cycle (**Figure 3C, 3D; Movie S5**). All remaining VPCs exited into a CDK^low^ state at their terminal division. Consistent with these results, bootstrap analyses (**Figure S2G-M**) support our qualitative results, such that we can quantitatively distinguish between a cell that has completed mitosis and will continue to cycle (CDK^inc^) from a cell that exits mitosis and enters a terminally differentiated G0 state (CDK^low^).

**Figure 3.**
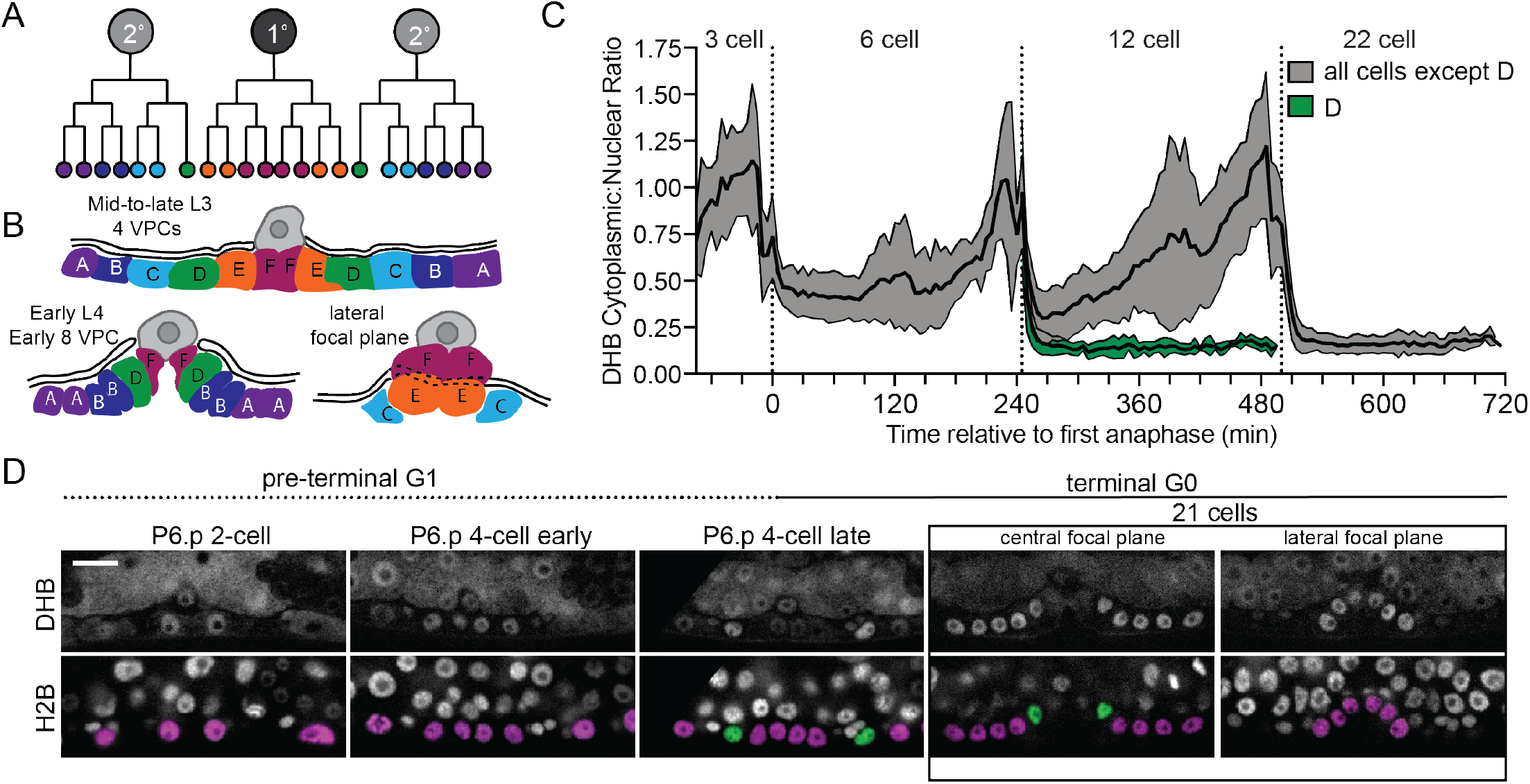
Vulval precursor cells exit terminal divisions into a CDK^low^ state. (A) Schematic of primary (1°) and secondary (2°) fated vulval precursor cells (VPCs). (B) All of the VPCs divide, with the exception of the D cells, to facilitate vulval morphogenesis. (C) Time series of CDK sensor localization in the 1° and 2° VPCs, as measured every 5 min. Note that the terminally differentiated D cells are born into a CDK^low^ state (*n*≥9 cells). Dotted line indicates time of anaphase. Shaded error bands depict mean±SD. (D) Representative images of CDK sensor localization in the VPCs from the P6.p 2-cell stage to 8-cell stage. Nuclei (H2B) are highlighted in magenta for non-D cell 1° and 2° VPCs and green for the D cells. Scale bar = 10 μm.

### CKI-1 Levels Peak Prior to Terminal Differentiation

In mammalian cell culture, endogenous levels of p21^Cip1^ during G2 are predictive of whether a cell will go on to divide or enter quiescence/senescence/terminal differentiation (Hsu et al., 2019; Moser et al., 2018; Overton et al., 2014; Spencer et al., 2013). This raises the intriguing possibility that endogenous levels of CKI-1 in *C. elegans* correlate with CDK^low^ or CDK^inc^ activity. To co-visualize CKI-1 dynamics with our CDK sensor, we inserted a N-terminal GFP tag into the endogenous locus of *cki-1* via CRISPR/Cas9 and introduced a DHB::2xmKate2 variant of the sensor (devoid of histone H2B) into this genetic background. Since endogenous levels of GFP::CKI-1 were too dim for time-lapse microscopy, likely due to its short half-life (Yang et al., 2017), we collected a developmental time series of static images over the L3 and L4 larval stages to characterize GFP::CKI-1 levels during pre-terminal and terminal divisions in the VPC lineage. We detected generally low levels of GFP::CKI-1 at the Pn.p 2-cell stage (**Figure 4A, 4B, S3A-C**). In their daughter cells, at the Pn.p 4-cell stage, we detected an increase in GFP::CKI-1 levels in cycling cells prior to their next cell division, peaking in G2 (**Figure 4A, 4C, S3A-C**). Notably, the D cell, which becomes post-mitotic after this cell division, exits mitosis with higher levels of GFP::CKI-1 than its CD mother (**Figure 4A, S3B**). This trend holds true for the remaining VPCs during terminal differentiation at the Pn.p 6-cell and 8-cell stage, which show high levels of GFP::CKI-1 that peak immediately after mitotic exit and remain high during the post-mitotic L4 stage (**Figure 4A, 4D, 4E, S4A-C**). We also observed increasing levels of GFP::CKI-1 in the G2 phase of mother cells that peak in their terminal daughter cells in the uterine (**Figure S3D**) and SM cell lineages (**Figure S3E**). Thus, levels of GFP::CKI-1 increase in mother cells prior to terminal differentiation and remain high upon mitotic exit in daughter cells with CDK^low^ activity. These results elucidate a proliferation-terminal differentiation decision process that is already underway in G2 of the previous cell cycle and is in part controlled by CKI-1 in the mother cell.

**Figure 4.**
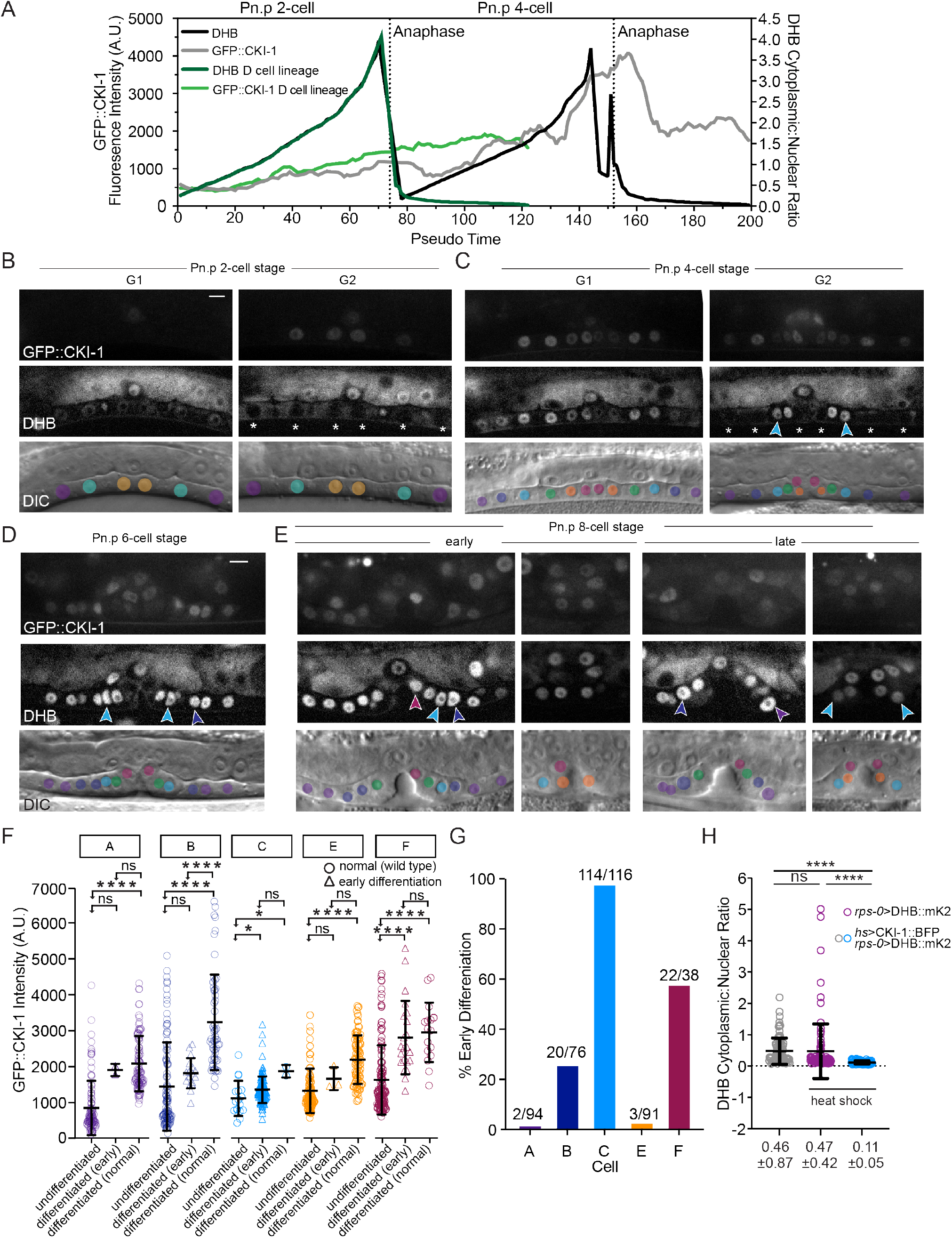
CKI-1 levels peak prior to terminal differentiation. (A) CDK sensor activity and CKI-1 levels across pseudo-time and DHB ratios for all VPCs (black line) and D cells (dark green line). GFP::CKI-1 fluorescence in VPCs (grey line) and D cell (light green line); *n*≥93 cells per lineage. (B) Representative images of VPCs at the Pn.p 2-cell stage at G1 and G2 (white asterisk). (C) Representative images of VPCs at the Pn.p 4-cell stage at G1 and G2; early differentiated C cells (cyan arrows) with low levels of GFP::CKI-1. (D, E) Representative images of VPCs at the Pn.p 6-cell stage (D) and 8-cell stage (E); arrows show early differentiated C (cyan) and B cell (dark blue), F cell (magenta), and A cell (purple). (F) GFP::CKI-1 fluorescence in each cell of the VPC lineage (*n*≥16, except C normal and A early *n*=2, E early *n*=3). (G) Percentage of cells of each lineage that showed signs of early differentiation and did not undergo their final division. (H) Overexpression of CKI-1 via heat shock causes pre-terminally differentiated cells to enter G0 (*n*≥36 cells per treatment). Scale bar = 10 μm. ns, not significant, \**p*≤0.05, \*\*\*\**p*≤0.0001. Significance determined by statistical simulations; *p*-values in Table S1.

During our collection of static images of GFP::CKI-1 animals, we observed significant deviations in the expected VPC lineage pattern in the early L4 larval stage. In particular, we noted that many cells appeared to bypass their final division and undergo early terminal differentiation with coincident high levels of GFP::CKI-1 and low DHB ratios. We hypothesized that the line we generated could be behaving as a gain-of-function mutant, as GFP insertions at the N-terminus could interfere with proteasome-mediated protein degradation of CKI-1 (Bloom et al., 2003). The penetrance of this early terminal differentiation defect varied across VPC lineages. While the A (2% of cases observed) and E (3% of cases observed) lineages showed a low penetrance of this early terminal differentiation defect, the B (26% of cases observed) and F (58% of cases observed) lineages showed a moderate penetrance (**Figure 4F, 4G**). We speculate that the A and E lineages are largely insensitive to the gain-of-function mutant because CKI-2, an understudied paralog of CKI-1, may be the dominant CKI in these cells. The C cell, sister to the terminally differentiated D cell, had a highly penetrant early terminal differentiation defect (98% of cases observed; **Figure 4F, 4G**). Consistent with our finding that high levels of endogenous GFP::CKI-1 can lead to early terminal differentiation, heat shock-induced CKI-1 expression uniformly drove VPCs into a CDK^low^ G0 state with mean DHB ratios of 0.11±0.05 (**Figure 4H**) as compared to control animals that lacked heat shock-induced expression (0.46±0.87) or lacked the inducible *cki-1* transgene (0.47±0.42). Together, these results demonstrate that cycling cells are highly sensitive to levels of CKIs and that increased expression can induce a terminally differentiated G0 state.

### CDK Activity Predicts a Cryptic Stochastic Fate Decision in an Invariant Cell Lineage

A strength of *C. elegans* is the organism’s robust ability to buffer external and internal perturbations to maintain its invariant cell lineage. However, not all cell divisions that give rise to the 959 somatic cells are completely invariant. Studies have identified several lineages, including the vulva, where environmental stressors, genetic mutations and/or genetic divergence of wild isolates leads to stochastic changes in a highly invariant cell fate pattern (Braendle and Felix, 2008; Hintze et al., 2020; Katsanos et al., 2017). Thus, we wondered if the CDK sensor generated here could be utilized to visualize and predict stochastic lineage decisions during *C. elegans* development.

The VPC lineage that gives rise to the adult vulva is invariant (**Figure 5A, S4A**) (Sulston and Horvitz, 1977). However, at high temperatures it has been observed that the D cell, the inner-most granddaughter of P5.p or P7.p, will go on to divide (**Figure 5A**) (Sternberg, 1984; Sternberg and Horvitz, 1986). Unexpectedly, we noticed a rare occurrence of D cells expressing elevated DHB ratios during the course of time-lapse analysis of VPC divisions captured under standard laboratory conditions. To determine the penetrance of the cycling D cell phenotype, we inspected each of our CDK sensor lines grown at 25°C, a high temperature that is still within normal range for *C. elegans*. In both strains we observed a cycling D cell with a 2-10% penetrance (**Figure 5B, S4B**). To test whether this cycling D cell phenotype resulted from the presence of the DHB transgene or environmental stressors, such as temperature fluctuation, we examined the VPC lineage in animals lacking the CDK sensor at 25°C and 28°C. At 25°C, we observed a low penetrance (2%) of cycling D cells in a strain expressing an endogenously tagged DNA licensing factor, CDT-1::GFP (**Figure 5B, S4B**), which is cytosolic in cycling cells (Matus et al., 2014; Matus et al., 2015). From lineage analysis, L2 larvae, expressing a seam cell reporter (*scm*>GFP), that were temperature shifted from 20°C to 28°C displayed approximately a 30% occurrence of extra D cell divisions (**Figure S4C-E**). Lastly, we wanted to determine whether D cells that show CDK^inc^ activity divide. To accomplish this, we collected time-lapses of DHB::GFP animals grown at 25°C. These time-lapses revealed 10 occurrences of D cells born into a CDK^inc^ rather than a CDK^low^ (**Figure 5C, 5D; Movie S6**). In all 10 cases, the CDK^inc^ D cell goes on to divide (**Figure S4A**). Thus, we find that CDK activity shortly after mitosis is a strong predictor of future cell behavior, even in rare stochastic cases of extra cell divisions in *C. elegans*, an organism with a well-defined cell lineage.

**Figure 5.**
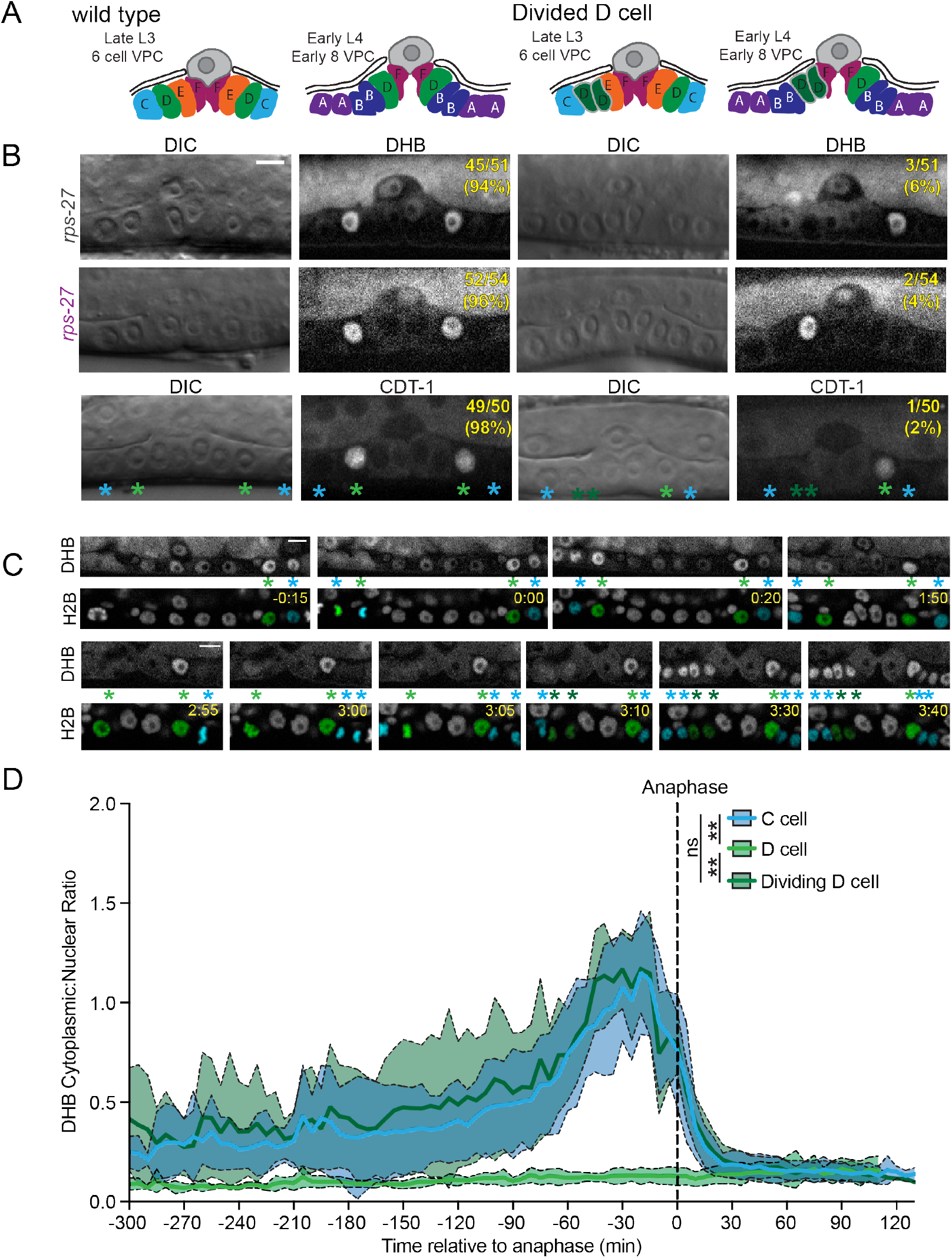
CDK activity predicts a cryptic stochastic fate decision in an invariant cell lineage. (A) Schematic of wild type vulva and vulva with a divided D cell. (B) Representative images at the Pn.p 6-cell stage from CDK sensor strains (top, middle) and endogenous *cdt-1::GFP* (bottom), showing wild type vulva on the left and vulva with a divided D cell on the right. Penetrance of each phenotype for each strain is annotated on the DHB image. (C) Frames from a time-lapse with a dividing D cell (left; see Movie S5). Nuclei (H2B) are highlighted in green for the D cell and cyan for the C cells. Green asterisks mark the D cell and cyan asterisks mark the C cell. Scale bar = 10 μm. (D) DHB ratio for C cell, terminally differentiated D cell and dividing D cell (D *n*=10 cells, C *n*=20 cell divisions and Dividing D *n*=10 cell divisions). Dotted line indicates time of anaphase. Line and shaded error bands depict mean±SD. ns, not significant, \*\**p*≤0.01. Significance determined by statistical simulations; *p*-values in Table S1.

### Generation of Inducible CDK Sensor Transgenic Lines in Zebrafish

To investigate the predictive capability of DHB ratios in zebrafish, we generated two CDK sensor lines with different fluorescent protein combinations, DHB-mNeonGreen (DHB-mNG) and DHB-mScarlet (DHB-mSc) with H2B-mSc and H2B-miRFP670, respectively, to allow for flexibility with imaging and experimental design (**Figure 6A**). Both transgenes are under the control of the *hsp70l* heat shock-inducible promoter, which produces robust ubiquitous expression after shifting the temperature from 28.5°C to 40°C for 30 min (Halloran et al., 2000; Shoji et al., 1998). We also generated a transgenic line, *Tg(ubb:Lck-mNG)*, that ubiquitously labels the plasma membrane with mNG, which we crossed into the HS:DHB-mSc-2A-H2B-miRFP670 line to simultaneously visualize CDK activity (DHB-mSc), segment nuclei (H2B-miRFP670) and segment the plasma membrane (LCK-mNG) (**Figure 6A**).

**Figure 6.**
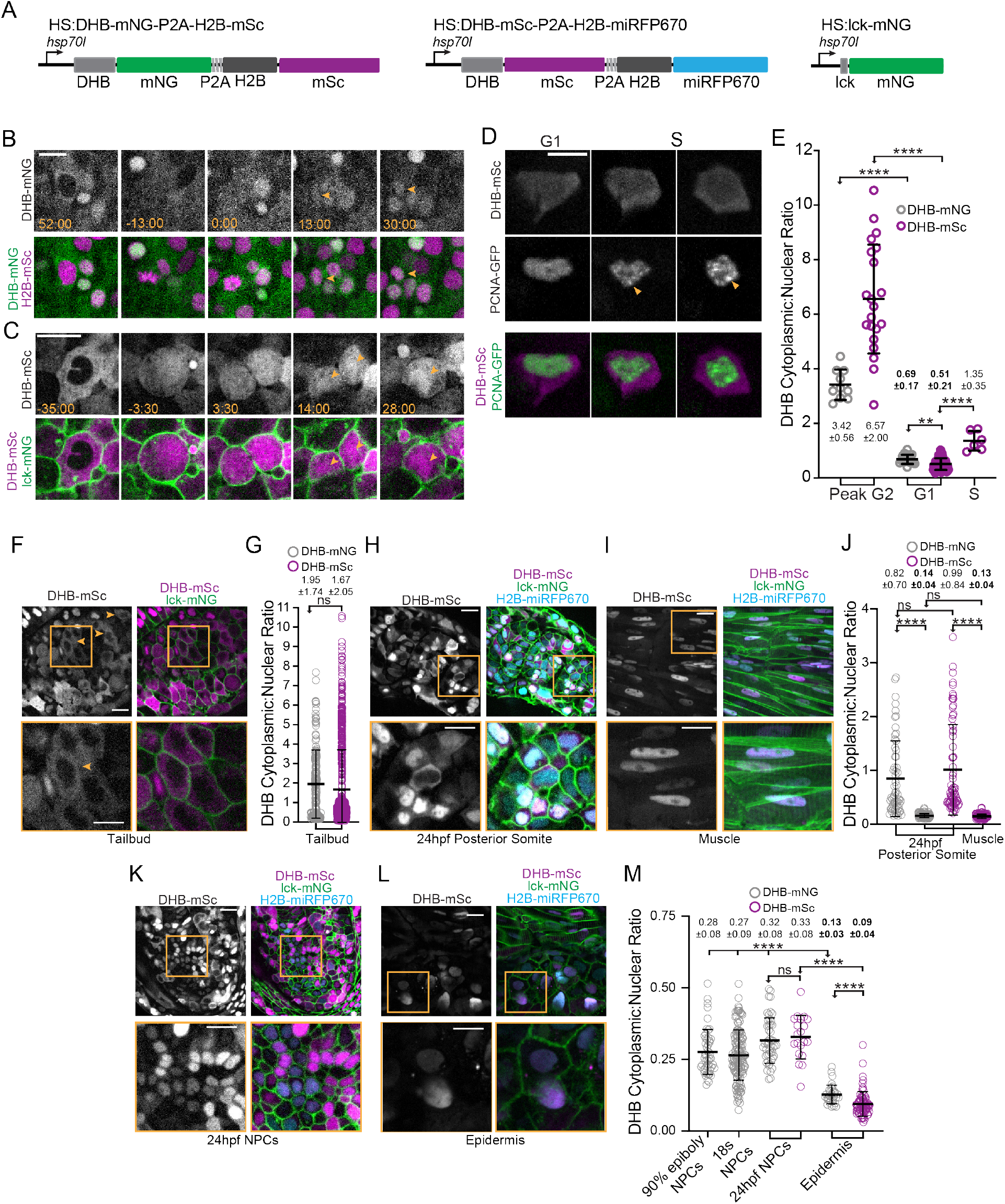
Generation of inducible CDK sensor transgenic lines in the zebrafish. Schematics of inducible zebrafish variants of the CDK sensor fused to mNG (left) or mSc (middle) and a nuclear mask (H2B-FP) separated by a self-cleaving peptide (P2A). Schematic of inducible membrane marker (lck-mNG; right). (B, C) Frames of DHB time-lapses. (D) DHB-mSC and PCNA-GFP puncta during S phase. (E) Dot plot of DHB ratios during interphase states (*n*≥7 cells from ≥2 embryos. (F) Representative micrographs of CDK sensor (orange arrows and box inset highlights cytosolic CDK sensor localization) and quantification of DHB ratio (G) in the tailbud (*n*≥160 cells). (H-M) Representative micrographs (H, I, K, L) and quantification of DHB ratios (J, M) in cells of 24 hpf posterior somites (*n*≥59 cells) (H) and terminally differentiated muscle at 72 hpf (*n*≥101) (I), notochord progenitors (NPCs) (*n*≥24 cells)(K) and epidermis at 72 hpf (*n*≥32 cells) (L). Insets, orange box, are zoom-ins. Scale bar = 20 μm. Line and error bars depict mean±SD. Numbers in **bold** are tissues in G0. ns, not significant, \*\**p*≤0.01, \*\*\*\**p*≤0.0001. Significance determined by statistical simulations; *p*-values in Table S1.

To verify that both CDK sensor lines localize DHB in a cell cycle-dependent manner, we first used time-lapse microscopy and quantified DHB ratios across cell divisions in the tailbud of bud or 22 somite-stage embryos (**Figure 6B, 6C, S5A**). We observed the expected localization pattern for both CDK sensor lines, with maximal nuclear exclusion of the sensor shortly before mitosis in G2 (3.42±0.56 (mNG) and 6.57±2.00 (mSc)) and low DHB ratios (0.69±0.17 (mNG) and 0.51±0.21 (mSC)) representing nuclear accumulation of the sensor shortly after mitosis in G1 (**Figure 6E, S5A**). To establish the DHB ratio for S phase we visualized PCNA-GFP in the tailbud of DHB-mSC embryos as PCNA forms puncta at S phase entry and returns to an even nuclear distribution in G2 (**Figure 6D**) (Leonhardt et al., 2000; Leung et al., 2011). Approximately 38.5 min after puncta formation, corresponding to mid S phase, the DHB ratio is 1.36±0.36, which is significantly higher than the G1 DHB value (0.51±0.21) (**Figure 6E**). Thus, we conclude that both CDK sensor lines localize in a cell cycle-dependent fashion, and that quantitative measurements can be used to determine interphase states.

Next, using both DHB transgenic lines, we examined CDK activity in a number of defined embryonic tissues. Imaging of the developing tailbud reveled cells in all phases of the cell cycle with a mean DHB ratios of 1.95±1.74 (mNG) and 1.67±2.05 (mSC) (**Figure 6E, 6F, S5B**). The tailbud of vertebrate embryos contain neuromesodermal progenitors (NMPs) (Martin, 2016), which in zebrafish have been reported to be predominantly arrested in the G2 phase of the cell cycle (Bouldin et al., 2014). Consistent with this, we observed cells with high CDK activity in the tailbud (orange arrows; **Figure 6F, S5B**). This enrichment is eliminated when embryos are treated with the CDK4/6 inhibitor palbociclib, leading to a significant increase of cells in the tailbud with low CDK activity (0.58±0.3), similar in range to the G1/G0 values we measured during time-lapse (0.69±0.17; **Figure S5C-E**). We also made the surprising observation that primitive red blood cells in the intermediate cell mass of 24 hours post-fertilization (hpf) embryos, which are nucleated in zebrafish, display high CDK activity (3.00±0.97) indicating that they are exclusively in the G2 phase of the cell cycle (**Figure S5F, S5G**), suggesting that cell cycle regulation may be important for hematopoiesis (Bronnimann et al., 2018; De La Garza et al., 2019)

### Visualization of Proliferation and Terminal Differentiation during Zebrafish Development

To examine differences between proliferating and terminally differentiated cells, we examined CDK activity in the somites, which are segmental mesodermal structures that give rise to terminally differentiated skeletal muscle cells and other cell types (Martin, 2016). In the most recently formed somites at 24 hpf, cells can be observed in all phases of the cell cycle (**Figure 6H, 6J, S5H**). Consistent with what we observed in the tailbud, treatment with palbociclib also caused somite cells to arrest with low CDK activity in G1/G0 (0.33±0.42; **Figure S5I-K**). Recently formed somites contain a subpopulation of cells called adaxial cells, which are positioned at the medial edge of the somite next to the axial mesoderm (**Figure S5L**). These are the slow muscle precursors and are considered to be in a terminally differentiated state through the cooperative action of Cdkn1ca (p57) and Myod (Osborn et al., 2011). As opposed to the majority of cells in the lateral regions of recently formed somites, adaxial cells possess low CDK activity (0.13±0.04 (mNG); **Figure S5M**). At later stages, the majority of cells in the lateral regions of the somite will differentiate into fast skeletal muscles fibers, which are also considered to be in a terminally differentiated state (Halevy et al., 1995). Examination of DHB ratios at 72 hpf skeletal muscle fibers revealed they have low CDK activity (0.14±0.04 (mNG) and 0.13±0.04 (mSc)), similar to the adaxial cells, but significantly different than the mean DHB ratios of undifferentiated cells at 24 hours (0.82±0.70 (mNG) and 0.99±0.084 (mSc); **Figure 6I, 6J, S5H**). Thus, from our static imaging, we can identify cell types with low CDK activity that are thought to be terminally differentiated.

We next sought to determine if we can differentiate between the G1 and G0 state based on ratiometric quantification of DHB. We compared the terminally differentiated muscle and epidermal cells to notochord progenitor cells, which are held transiently in G1/G0 before re-entering the cell cycle upon joining the notochord (Sugiyama et al., 2014; Sugiyama et al., 2009) (**Figure 6I**). Notably, the mean DHB-mNG ratio of the notochord progenitors (0.32±0.08) is significantly higher than the DHB-mNG ratio of the differentiated epidermis (0.13±0.03; **Figure 6K-M, S5N, S5O**). This value is consistent in notochord progenitors at two other earlier developmental stages, 90% epiboly (0.28±0.08) and 18 somites (0.27±0.09; **Figure 6M**). The DHB-mSc ratio in the notochord progenitors (0.33±0.08) is also significantly different than the differentiated epidermis (0.09±0.04; **Figure 6M**). Based on this difference in DHB ratios between notochord progenitors and terminally differentiated cell types, including muscle (**Figure 6I, 6J**) and epidermis (**Figure 6L, 6M**), and our knowledge of the normal biology of these cells, we conclude that the CDK sensor can distinguish between a cycling G1 state and a terminally differentiated G0 state in the zebrafish.

### A Bifurcation in CDK Activity at Mitosis is Conserved in *C. elegans* and Zebrafish

We next investigated whether zebrafish cells separate into G1/CDK^inc^ and G0/CDK^low^ populations as they do in the nematode *C. elegans* and whether these CDK activity states are a general predictor of future cell behavior in both animals. First, we plotted all of the time-lapse CDK sensor data we collected in *C. elegans* (**Figure 7A, 7B**) and zebrafish (**Figure 7C, 7D**). For *C. elegans*, plotting of all CDK sensor trace data, irrespective of lineage, demonstrated that cells entering a CDK^low^ state after mitosis corresponded to terminally differentiated cells, while cells that exited mitosis into a CDK^inc^ state corresponded to cells from pre-terminal divisions. For zebrafish, in a lineage agnostic manner, we plotted all the traces from the tailbud and found that these data could also be classified into CDK^low^ and CDK^inc^ populations.

**Figure 7.**
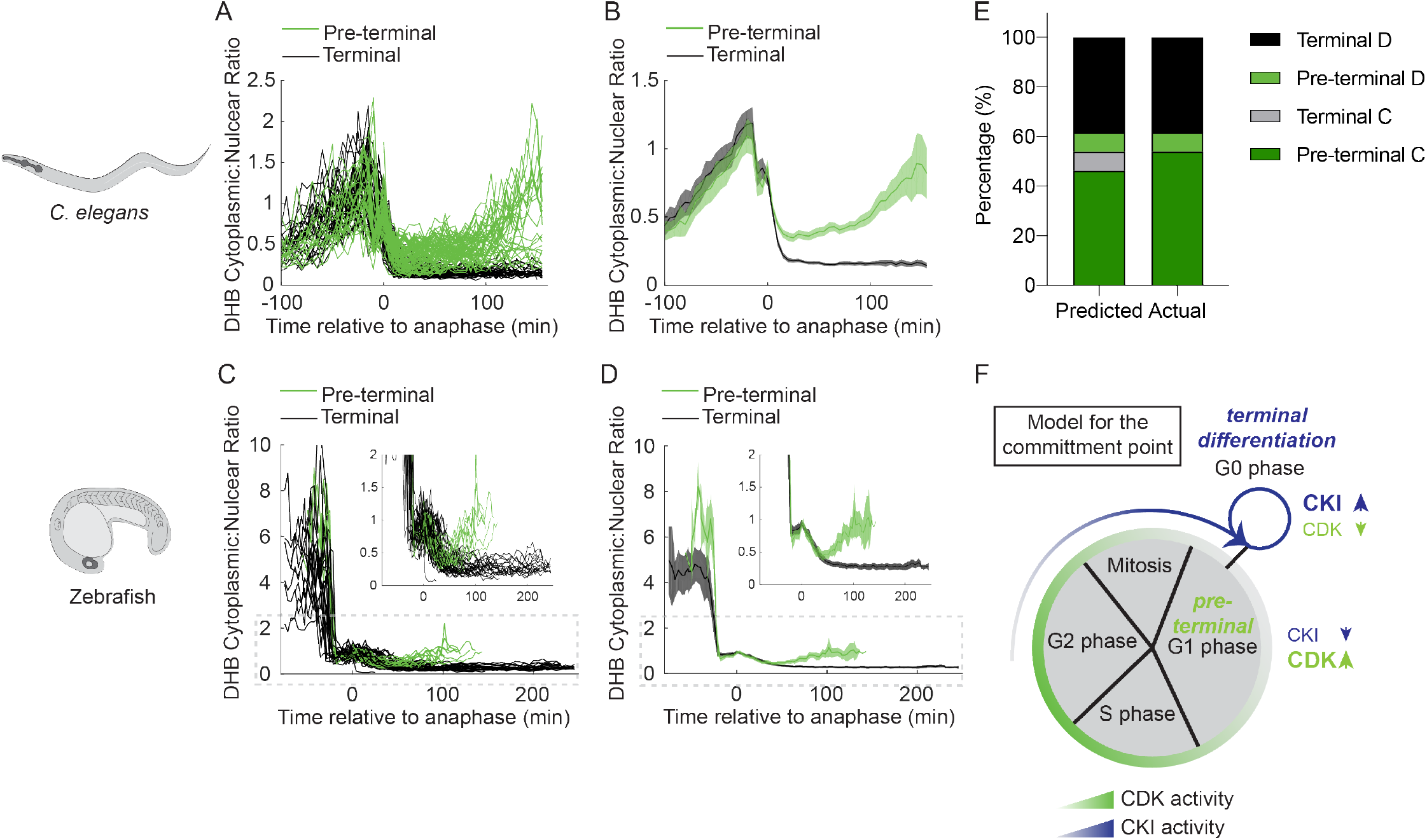
A bifurcation in CDK activity at mitotic exit controls the proliferation-differentiation decision. (A-D) Single-cell traces of CDK activity for all quantified *C. elegans* (A-B) and zebrafish (C-D) cell births for CDKinc cells (green) and CDK^low^ cells (black). DHB ratio of single-cell data (A, C) and mean±95% confidence interval (B, D) are plotted for each cell analyzed relative to anaphase. (E) A model for the metazoan commitment point argues that the G1/G0 decision is influenced by a maternal input of CKI activity and that CDK activity shortly after mitotic exit determines future cell fate.

As we were able to detect a rare stochastic lineage change in the *C. elegans* vulval lineage (**Figure 5**), we selected all CDK sensor trace data from the *C. elegans* VPCs (**Figure S6A**) and used this data to build a classifier to predict pre-terminal (G1) versus terminal (G0) cell fates based on CDK activity after anaphase (**Figure S6A-C**). Cross-examining our modeling with the known cell VPC lineage demonstrated that at 20 min after anaphase we had 85% accuracy in predictions with near perfect prediction 60 min after anaphase (**Figure S6B**). To test the predictive power of the classifier, we analyzed CDK trace data from the births of C and D cells, where some D cells stochastically divide (**Figure 5**). Our classifier correctly predicted cell fate 92% (*n*=24/26 single cell traces) of the time, including the two occurrences of a stochastic mitotic D cell in the data set (**Figure 7E**). Together, these results demonstrate that during development, cycling cells encounter a bifurcation in CDK activity following mitosis where they either: (1) increase in CDK activity and become poised to cycle, or (2) exit into a CDK^low^ state and undergo terminal differentiation (**Figure 7F**). Thus, we suggest a model where cells from developing tissue in *C. elegans* and zebrafish must cross an early commitment point in the cell cycle where these cells must make the decision to divide or terminally differentiate. The decision to undergo terminal differentiation is crucial to tissue integration and organization and is controlled by the activity of evolutionarily conserved CKI(s) in the mother cell (**Figure 7F**) that control daughter cell CDK activity.

## DISCUSSION

### A CDK Sensor for Live-Cell In Vivo Imaging of Interphase States and the G1/G0 Transition

We introduce here a CDK activity sensor to visually monitor interphase and the proliferation-terminal differentiation decision in real-time and *in vivo* in two widely used research organisms, *C. elegans* and zebrafish. This sensor, which reads out the phosphorylation of a DHB peptide by CDKs (Hahn et al., 2009; Spencer et al., 2013), allows for quantitative assessment of cell cycle state, including G0. The use of FUCCI in zebrafish (Bouldin and Kimelman, 2014; Sugiyama et al., 2009) and past iterations of a CDK sensor in *C. elegans* (Deng et al., 2020; Dwivedi et al., 2019; van Rijnberk et al., 2017) and *Drosophila* (Hur et al., 2020) have been informative in improving our understanding of cell cycle regulation of development, but have not addressed the proliferation-terminal differentiation decision. The DHB transgenic lines generated in this study will allow researchers to distinguish G1 from G0 shortly after a cell has divided and directly study G0-related cell behaviors, such as terminal differentiation, quiescence and senescence, in living organisms.

Previously, CDK sensors have been used to distinguish between proliferative and quiescent cells in asynchronous mammalian cell culture populations (Arora et al., 2017; Cappell et al., 2016; Gast et al., 2018; Gookin et al., 2017; Miller et al., 2018; Moser et al., 2018; Overton et al., 2014; Spencer et al., 2013; Yang et al., 2015). As mammalian cells complete mitosis, they are born into either a CDK2^inc^ state in which they are more likely to divide again or a CDK2^low^ state in which they remain quiescent. Here we have examined the CDK activity state of cells in an invertebrate with a well-defined and invariant lineage, *C. elegans*, and a vertebrate that lacks a defined cell lineage, the zebrafish. In both contexts, we can visually and quantitatively differentiate between cells that are in a CDK^inc^ state following cell division and cells that are in a CDK^low^ state. Strikingly, in *C. elegans* these states precisely correlate with the lineage pattern of the three post-embryonic tissues we examined: the SM cells, uterine cells, and VPCs. Cells born into a CDK^inc^ state represented pre-terminal divisions, whereas cells born into a CDK^low^ state were terminally differentiated. By distinguishing these two states in CDK activity, we were able to accurately identify shortly after cell birth a rare stochasticity that was first described through careful end-point lineage analysis nearly 36 years ago in the *C. elegans* vulval lineage (Sternberg, 1984; Sternberg and Horvitz, 1986). Further, statistical modeling demonstrated that we could predict future cell behavior with >85% accuracy in *C. elegans* just 20 min post-anaphase. In zebrafish, we found that we could readily distinguish between CDK^inc^ cells in G1, such as notochord progenitors, which re-enter the cell cycle after joining the notochord, and tissues that are terminally differentiated and contain CDK^low^ cells in G0, such as skeletal muscle and epidermis. Thus, in both organisms the CDK sensor can be easily used to separate G1 from G0 without the need for multiple fluorescent reporters (Bajar et al., 2016; Oki et al., 2014) or fixation followed by antibody staining for FACS analysis (Tomura et al., 2013).

### In Vivo Evidence of a G2 Commitment Point in the Metazoan Cell Cycle

The classic model of the Restriction Point, the point in G1 at which cells in culture decide to commit to the cell cycle and no longer require growth factors (e.g., mitogens), is that mammalian cells are born uncommitted and that the cell cycle progression decision is not made until several hours after mitosis (Jones and Kazlauskas, 2001; Pardee, 1974; Zetterberg and Larsson, 1985; Zwang et al., 2011). An alternative model has been proposed in studies using single-cell measurements of CDK2 activity in asynchronous populations of MCF10A cells (Spencer et al., 2013) and other nontumorigenic as well as tumorigenic cell lines (Moser et al., 2018). This model extends the classic Restriction Point model for cell cycle commitment. During the G2 phase of the cell cycle, the mother cell is influenced by levels of p21 and cyclin D and these levels affect the phosphorylation state of Rb in CDK^low^ and CDK^inc^ daughter cells, respectively (Min et al., 2020; Moser et al., 2018). In CDK^low^ daughter cells, phospho-Rb is low and these cells are still sensitive to mitogens. Whether cells *in vivo* coordinate cell cycle commitment with levels of CKI and CDK over this extended Restriction Point was poorly understood.

By first quantifying the cytoplasmic:nuclear ratio of the CDK sensor in time-lapse recordings of cell divisions in *C. elegans* somatic lineages, we were able to use DHB ratios as a proxy for CDK levels to distinguish two populations of daughter cells: the first being actively cycling cells in a CDK^inc^ state (G1), and the second being terminally differentiated cells in a CDK^low^ state (G0). We then quantified cytoplasmic:nuclear ratio of the CDK sensor in time-lapse recordings of cell divisions in zebrafish and we were also able to distinguish two populations of daughter cells. As data from asynchronous cell culture studies suggest that the decision to commit to the cell cycle is made by the mother cell as early as G2 (Moser et al., 2018; Spencer et al., 2013), we wanted to determine if this same phenomenon occurred *in vivo*. To accomplish this, we endogenously tagged one of two CKIs in the *C. elegans* genome, *cki-1*, with GFP using CRISPR/Cas9. We paired static live-cell imaging of GFP::CKI-1 with DHB::2xmKate2 during vulval development. Similar to *in vitro* experiments (Moser et al., 2018; Spencer et al., 2013), we found that mother cells whose daughters are born into a CDK^inc^ G1 state will divide again, expressing low levels of GFP::CKI-1. In contrast, mother cells of daughters that will differentiate express a peak of GFP::CKI-1 in G2 which increases as daughter cells are born into a CDK^low^ G0 state. Thus, our data demonstrate that an extended Restriction Point exists in the cell cycle of intact Metazoa, and that the *in vivo* proliferation-terminal differentiation decision can be predicted in *C. elegans* by CKI/CDK activity shortly after mitotic exit.

## Conclusion

We demonstrate here that the CDK sensor functions in both *C. elegans* and zebrafish to read out cell cycle state dynamically, and unlike other cell cycle sensors, can distinguish between proliferative and terminally differentiated cells within an hour of cell birth. As nematodes and vertebrates last shared a common ancestor over 500 million years ago, this suggests that the CDK sensor is likely to function in a similar fashion across Metazoa. The broad functionality of the sensor offers researchers a unique opportunity to examine fundamental questions such as the relationship between cell cycle state and cell fate during normal development, cellular reprogramming, and tissue regeneration. Finally, as an increasing body of evidence suggests that cell cycle state impinges on morphogenetic events ranging from gastrulation (Grosshans and Wieschaus, 2000; Murakami et al., 2004), convergent extension (Leise and Mueller, 2004) and cellular invasion (Kohrman and Matus, 2017; Matus et al., 2015; Medwig-Kinney et al., 2020), this CDK sensor will provide the means to increase our understanding of the relationship between interphase states and morphogenesis during normal development and diseases arising from cell cycle defects, such as cancer.

## SUPPLEMENTAL INFORMATION

Supplemental Information includes Extended Experimental Procedures, six figures, one table, and seven movies, which can be found online with this article.

## ACKNOWLEDGEMENTS

We thank D. Özpolat, D. Pisconti, C.-K. Hu and M.J. Gacha-Garay for helpful comments on the manuscript; M.J. Gacha-Garay, N. Bhattacharji and S. Flanagan for fish care; J. Maghakian and L. Yang for consultation on statistical modeling; D. Dickinson and B. Goldstein for assistance with the CRISPR single copy knock-in strategy; and T. Geer of Nobska Imaging, Inc. for helping maintain our spinning disk confocal microscopes. This work was funded by the NIH NIGMS [1R01GM121597-01 to D.Q.M. and 1R01GM124282 to B.L.M.]. D.Q.M. and B.L.M. are both Damon Runyon-Rachleff Innovators supported by the Damon Runyon Cancer Research Foundation [DRR-47-17]. B.L.M. also received support from the NSF [IOS 1452928] and the Pershing Square Sohn Cancer Research Alliance. R.C.A., A.Q.K., J.J.S. and M.A.Q.M. are all supported by the NIGMS [1F32133131-01, F31GM128319-01, 3R01GM121597-02S1/S2, respectively]. T.N.M-K. is supported by the NIH NICHD [F31HD100091-01]. N.J.P. is supported by the ACS [132969-PF-18-226-01-CSM]. J.L.F. and S.L.S. are both supported by an NIH Director’s New Innovator Award (DP2GM1191136-01 and DP2-CA238330, respectively). S.L.S. is also supported by an ACS Research Scholar Grant (RSG-18-008-01), a Pew-Stewart Scholar Award, a Beckman Young Investigator Award, a Boettcher Webb-Waring Early-Career Investigator Award, a Kimmel Scholar Award (SKF16-126), and a Searle Scholar Award (SSP-2016-1533). Some strains were provided by the *Caenorhabditis* Genetics Center, which is funded by the NIH ORIP [P40 OD010440].

## AUTHOR CONTRIBUTIONS

Conceptualization, R.C.A., A.Q.K., M.A.Q.M., M.D.S., J.L.F., S.L.S., B.L.M. and D.Q.M.; Methodology, R.C.A., A.Q.K., M.A.Q.M., N.J.P., J.J.S., T.N.M-K., S.L., R.D.M., W.Z., B.L.M., and D.Q.M.; Formal Analysis, R.C.A., M.A.Q.M., N.J.P., A.Q.K., T.N.M-K., M.M., S.L.S, B.L.M. and D.Q.M.; Investigation, R.C.A., A.Q.K., M.A.Q.M., N.J.P., J.J.S., M.D.S., O.B.A., N.K., N.W., M.B., A.M.P., B.L.M. and D.Q.M.; Writing, M.A.Q.M, R.C.A., A.Q.K., T.N.M-K., B.L.M and D.Q.M.; Visualization, R.C.A., M.A.Q.M, T.N.M-K. and D.Q.M.; Funding Acquisition, A.Q.K., R.C.A., N.J.P., T.N.M-K., B.L.M. and D.Q.M.

## EXPERIMENTAL PROCEDURES

### *C. elegans* Strains

*C. elegans* strains were cultured in standard conditions at 15-25°C on NGM plates with *E. coli* OP50. In the text and figures, we designate linkage to a promoter with a greater than symbol (>) and use a double colon (::) for linkages that fuse open reading frames (Ziel et al., 2009). See Extended Experimental Procedures and the Key Resources Table for details about alleles and transgenes generated in this study.

### Zebrafish Lines

All zebrafish experiments and husbandry were performed with approval from the Stony Brook University Institutional Animal Care and Use Committee. See Extended Experimental Procedures and the Key Resources Table for details about the transgenic lines generated in this study.

### RNAi, Heat Shock Induction and Chemical Inhibitors

RNAi was delivered by feeding worms *E. coli* HT115(DE3) expressing double-stranded RNA (dsRNA) targeted against *control* (L4440) and *cdk-1*. Expression of *cki-1* was induced using a ubiquitous heat shock promoter (**Figure 2G, 4H**). The inhibitor used in zebrafish was PD-0332991, a CDK4/6 inhibitor. See Extended Experimental Procedures for details.

### Live-Cell Microscopy

For microscopes and imaging conditions, see Extended Experimental Procedures.

### Image Processing and Statistical Analyses

Image processing was performed in Fiji and statistical analyses were performed in MATLAB. See Extended Experimental Procedures.

