## Supplemental Materials for "Visualizing the metazoan proliferation-terminal differentiation decision *in vivo*"

**Key Resources Table**

| REAGENT or RESOURCE | SOURCE | IDENTIFIER |
|---|---|---|
| ***Bacterial Strains*** | | |
| RNAi feeding strain | *Caenorhabditis* Genetics Center | *E. coli* HT115(DE3) |
| Ahringer RNAi Library | Source BioScience | *C. elegans* RNAi Collection |
| ***Chemicals*** | | |
| Carbenicillin | Alfa Aesar | J61949 |
| Isopropyl-β-D-thiogalactoside | Thermo Scientific | R0393 |
| Palbociclib | MedChemExpress | HY-A0065 |
| Hygromycin B | Millipore | 400052 |
| Sodium Azide | Sigma-Aldrich | S2002 |
| Tricaine (*C. elegans*) | Sigma-Aldrich | E10521 |
| Tricaine-S (MS-222) (zebrafish) | Pentair | TRS1 |
| Levamisole | Sigma-Aldrich | L9756 |
| ***C. elegans Strains*** | | |
| *LoxN::rps-27>DHB::GFP::P2A::H2B::2xmKate2 (bmd86) LGI* | This study | DQM298 |
| *LoxN::rps-0>DHB::mKate2 (bmd118) LGII* | This study | DQM394 |
| *LoxN::hsp16-41>cki-1::2xmTagBFP2 (bmd129) LGI; LoxN::rps-0>DHB::mKate2 (bmd118) LGII* | This study | DQM406 |

| | | |
|---|---|---|
| *LoxN::rps-27>DHB::2xmKate2::P2A::H2B::GFP (bmd147) LGI* | This study | DQM543 |
| *LoxN::pcn-1>PCN-1::GFP (bmd200) LGI; LoxN::rps-27>DHB::2xmKate2 (bmd168) LGII* | This study | DQM662 |
| *LoxN::rps-27>DHB::2xmKate2 (bmd156) LGI; GFP::LoxN::CKI-1::3xFLAG (bmd132) LGII* | This study | DQM586 |
| *cdt-1::ZF::LoxP::GFP::3xFLAG (wow98) LGI; zif-1(gk117) LG III** | This study | JLF634 |
| *SCM>GFP (wls51) LGV* | *Caenorhabditis* Genetics Center | JR667 |
| ***D. rerio Strains*** | | |
| *Tg(hsp70l:DHB.mNeonGreen-p2a-H2B.mScarlet)* | This study | Sbu108 |
| *Tg(ubb:Lck.mNeonGreen)* | This study | Sbu107 |
| *Tg(hsp70l:DHB.mScarlet-p2a-H2B.miRFP670)* | This study | Sbu109 |
| ***Recombinant DNA*** | | |
| *control RNAi* | Fire et al., 1998 | L4440; RRID: Addgene_1654 |
| *cdk-1 RNAi* | Kamath et al., 2003 | Ahringer RNAi library |
| *NotI-rps-27-NotI>DHB-ClaI-GFP-P2A-H2B-2xmKate2-NheI-3xHA (I)* | This study | pAWK61 |
| *NotI-rps-0>DHB-mKate2(GLO) - NheI-3xHA (II)* | This study | pAWK41 |
| *NotI-rps-27-NotI>DHB-2xmKate2-P2A-H2B-GFP-NheI-3xHA (I)* | This study | pWZ186 |
| *NotI-rps-27-NotI>DHB-2xmKate2 (I)* | This study | pWZ194 |
| *NotI-rps-27-NotI>DHB-2xmKate2 (II)* | This study | pTNM054 |
| *NotI-ccdB-ClaI-GFP-3xHA (I)* | This study | pWZ111 |
| *NotI-pcn-1>PCN-1-GFP-3xHA (I)* | This study | pWZ157 |
| *hsp-16.41-NotI-ccdB-ClaI-2xmTagBFP2-NheI-3xHA (I)* | This study | pWZ123 |

| | | |
|---|---|---|
| *hsp-16.41>CKI-1-2xmTagBFP2-NheI-3xHA (I)* | This study | pWZ199 |
| *ccdB-GFP-C1^SEC^3xFLAG-ccdB* | Dickinson et al., 2015 | pDD282 |
| *GFP-C1^SEC^3xFLAG-cki-1 (II)* | This study | pNJP026 |
| *cki-1 sgRNA* | This study | pWZ143 |
| *ccdB-ZF-GFP-SEC-3xFLAG-ccdB* | Sallee et al. 2018 | pJF250 |
| *cdt-1-ZF-GFP-3xFLAG (I)* | This study | pMS254 |
| *cdt-1 sgRNA* | This study | pMS250 |
| *hsp70I-DHB-mNeonGreen-P2A-h2b::mScarlet* | This study | pRM14 |
| *ubb-Lck-mNeonGreen* | This study | pRM27 |
| *hsp70I-DHB-mScarlet-P2A-h2b::miRFP670* | This study | pRM15 |
| *Hs-PCNA-GFP* | Strzyz et al., 2015 | Addgene:105942 |
| *sgRNA sequences* | | |
| gaagacatttgaaaagagtg | This study | *cki-1* sgRNA |
| ggatggccgtggtgtgtgg | This study | *cdt-1* sgRNA |
| ***Oligonucleotides*** | | |
| Primer: *rps*-27 F to insert in NotI site of pAP88 catcctgtaaaacgacggccagtgcTTCAATCGGTTTTTCCTTG | This study | DQM205 |
| Primer: *rps*-27 R to insert in NotI site of pAP88 ctctttttgacatacttcgggtagcggccgcTTTTATTCCACTTGTTGAGC | This study | DQM206 |
| Primer: *rps-0* F catcctgtaaaacgacggccagtgcGAGGAATGAAGAAATTTGC | This study | DQM728 |
| Primer: *rps-0* R cggaccaggtgacgtcgttggtcatATTACCTTAAAATTCAAAAATTAATTTCAG | This study | DQM729 |
| Primer: *pcn-1* F to insert into NotI-ccdB-ClaI site of pWZ111 CATCCtgtaaaacgacggccagtgcGGCCGCagaaacagtggccgtattgg | This study | DQM622 |
| Primer *pcn-1* R to insert into NotI-ccdB-ClaI site of pWZ111 tgaacaattcttctcctttactcatcgatgctccGTCCATATTCTCGTCGTC | This study | DQM609 |

| | | |
|---|---|---|
| Primer: *hsp* F catcctgtaaaacgacggccagtgcCACCAAAAACGGAACGTTGAGC | This study | DQM288 |
| Primer: *hsp* R ctctttttgacatacttcgggtagcggccgCCAATCCCGGGGATCCGA | This study | DQM289 |
| Primer: *cki-1* F atccccgggattggcggccgcATGTCTTCTGCTCGTCGTTG | This study | DQM303 |
| Primer: *cki-1* R aatcaattccgaaaccattgaggctcccgatgctccGTATGGAGAGCATGAAGATCG | This study | DQM304 |
| Primer: *gfp::cki-1* F atgttacccatccaactatacacc | This study | WZ1 |
| Primer: *gfp::cki-1* R gtggttctgacagtgagaac | This study | NP63R |
| Primer: cki-1 sgRNA tcctattgcgagatgtcttggaagacatttgaaaagagtgGTTTTAGAGCTAGAAATAGC | This study | DQM490 |
| Primer: *cdt-1* 5'HA (homology arm) F ttgtaaaacgacggccagtcg | This study | oMS-219-F |
| Primer: *cdt-1* 5'HA R CATCGATGCTCCTGAGGCTCC | This study | oMS-220-R |
| Primer: *cdt-1* 3'HA F CGTGATTACAAGGATGACGATGACAAGAGATAAaaactaatttctaagccatttgtaactaattttctcact | This study | oMS-208-F |
| Primer: *cdt-1* 3'HA R ggaaacagctatgaccatgttatcgatttcccaacgaggcgattactgagc | This study | oMS209-R |
| Primer: *cdt-1* sgRNA F GGATGGCCGTGGTGTGTGGgttttagagctagaaatagcaagt | This study | oMS-205-F |
| Primer: *cdt-1* sgRNA R CAAGACATCTCGCAATAGG | This study | oJF436-R |
| Primer: *mNG-Lck* F ATGGGCTGCGTGTGCAGCAGCAACCCCGAGATGGTGAGCAAGGGCGA | This study | RM112 |
| Primer: *mNG-Lck* R CTTGTACAGCTCGTCCATGC | This study | RM113 |

| | | |
|---|---|---|
| Primer: *mNG-Lck* homology F ATCTTACTTTGAATTTGTTTACAGGgatccATGGGCTGCGTGTGCAGCAG | This study | RM114 |
| Primer: *mNG-Lck* homology R TCATGTCTGGATCATCATCGATCTTGTACAGCTCGTCCATGCCC | This study | RM115 |
| Primer: *DHB:mNG* F ATCTTACTTTGAATTTGTTTACAGGgatccatgacaaatgatgtcacctggagc | This study | RM169 |
| Primer: *DHB:mNG* R GGTGGCGACCGGTGGAAC | This study | RM173 |
| Primer: *mScarlet:CAAX* F GTTCCACCGGTCGCCACCATGGTGAGCAAGGGCGAG | This study | RM174 |
| Primer: *mScarlet:CAAX* R CTTATCATGTCTGGATCATCATCGATCTTGTACAGCTCGTCCATGCC | This study | RM175 |
| Primer: *DHB* F AAGCTACTTGTTCTTTTTGCAGGATCCATGACAAATGATGTCACCTGGAGCGAG | This study | RM192 |
| Primer: *DHB* R GCCGCTGCCCTGGGCC | This study | RM193 |
| Primer: *mScarlet* F GCCCAGGGCAGCGGCATGGTGAGCAAGGGCGAG | This study | RM194 |
| Primer: *mScarlet* R GTTGGTGGCGCCGCTGCCCTTGTACAGCTCGTCCATGCC | This study | RM195 |
| Primer: *P2A:H2B* F GGCAGCGGCGCCACC | This study | RM196 |
| Primer: *P2A:H2B* R GGTGGCGACCGGTGGAACCT | This study | RM197 |
| Primer: *miRFP670* F AGGtTCCACCGGTCGCCACCATGGTAGCAGGTCATGCCTC | This study | RM198 |
| Primer: *miRFP670* R CTTATCATGTCTGGATCATCATCGATTTAGCTCTCAAGCGCGGTGA | This study | RM199 |
| ***Synthetic DNAs*** | | |

| | | |
|---|---|---|
| *Codon optimized DHB (with synthetic introns)* | catcctgtaaaacgacggccagtgcgg<br>ccgcATGACCAACGACGTCA<br>CCTGGTCCGAGGCCTCCTC<br>CCCAGACGAGCGTACCCTC<br>ACCTTCGCCGAGCGTTGGC<br>AACTCTCCTCCCCAGACGG<br>AGTCGACACCGACGACGA<br>CCTCCCAAAGTCCCGTGCC<br>TCCAAGCGTACCTGCGGAG<br>TCAACGACGACGAGTCCCC<br>ATCCAAGgtaagtttaaacatatata<br>tactaactaaccctgattatttaaattttca<br>gATCTTCATGGTCGGAGAG<br>TCCCCACAAGTCTCCTCCC<br>GTCTCCAAAACCTCCGTCT<br>CAACAACCTCATCCCACGT<br>CAACTCTTCAAGCCAACCG<br>ACAACCAAGAGACCGGAG<br>CATCGGGAGCCTCAGGAG<br>CATCGATGAGTAAAGGAGA<br>AGAATTGTTCA | Integrated DNA Technologies |
| *NgoMIV-P2A(codon-de-optimized)-his-58-GFP-NheI* | CATCCAAGCTCGGACATCG<br>TGCCGGCGCGGGAAGTGG<br>GGCCACGAACTTCAGTCTC<br>CTCAAACAAGCCGGGGAC<br>GTCGAAGAGAACCCCGGG<br>CCAATGCCACCAAAGCCAT<br>CTGCCAAGGGAGCCAAGA<br>AGGCCGCCAAGACCGTCG<br>TTGCCAAGCCAAAGGACGG<br>AAAGAAGAGACGTCATGCC<br>CGCAAGGAATCGTACTCCG<br>TCTACATCTACCGTGTTCTC<br>AAGCAAGTTCACCCAGACA<br>CCGGAGTCTCCTCCAAGGC<br>CATGTCTATCATGAACTCCT<br>TCGTCAACGATGTATTCGA<br>ACGCATCGCTTCGGAAGCT<br>TCCCGTCTTGCTCATTACA<br>ACAAACGCTCAACGATCTC<br>ATCCCGCGAAATTCAAACC<br>GCTGTCCGTTTGATTCTCC<br>CAGGAGAACTTGCCAAGCA<br>CGCCGTGTCTGAGGGAAC<br>CAAGGCCGTCACCAAGTAC<br>ACTTCCAGCAAGATGAGTA<br>AAGGAGAAGAATTGTTCAC<br>TGGAGTTGTCCCAATCCTC<br>GTCGAGCTCGACGGAGAC<br>GTCAACGGACACAAGTTCT | Twist Biosciences |

| | | |
|---|---|---|
| | CCGTCTCCGGAGAGGGAG<br>AGGGAGACGCCACCTACG<br>GAAAGCTCACCCTCAAGTT<br>CATCTGCACCACCGGAAAG<br>CTCCCAGTCCCATGGCCAA<br>CCCTCGTCACCACCTTCTG<br>CTACGGAGTCCAATGCTTC<br>TCCCGTTACCCAGACCACA<br>TGAAGCGTCACGACTTCTT<br>CAAGTCCGCCATGCCAGAG<br>GGATACGTCCAAGAGCGTA<br>CCATCTTCTTtAAGgtaagtttaa<br>acatatatatactaactactgattatttaa<br>attttcagGACGACGGAAACTA<br>CAAGACCCGTGCCGAGGT<br>CAAGTTCGAGGGAGACACC<br>CTCGTCAACCGTATCGAGC<br>TCCAGgtaagtttaaacagttcggta<br>ctaactaaccatacatatttaaattttcag<br>GGAATCGACTTCAAGGAGG<br>ACGGAAACATCCTCGGACA<br>CAAGCTCGAGTACAACTAC<br>AACTCCCACAACGTCTACA<br>TCATGGCCGACAAGCAAAA<br>GAACGGAATCAAGGTCAAC<br>TTCAAGgtaagtttaaacatgattta<br>ctaactaactaatctgatttaaattttcag<br>ATCCGTCACAACATCGAGG<br>ACGGATCCGTCCAACTCGC<br>CGACCACTACCAACAAAAC<br>ACCCCAATCGGAGACGGA<br>CCAGTCCTCCTCCCAGACA<br>ACCACTACCTCTCCACCCA<br>ATCCGCCCTCTCCAAGGAC<br>CCAAACGAGAAGCGTGACC<br>ACATGGTCCTCCTCGAGTT<br>CGTCACCGCCGCCGGAAT<br>CACCCACGGAATGGACGA<br>GCTCTACAAGTCAGGAGCT<br>AGCGGAGCCTACCCTTACG<br>ACG | |
| *gfp::cki-1* left homology arm | ACGTTGTAAAACGACGGCC<br>AGTCGCCGGCACTCACTGT<br>CACCAAATGTACCGTATTG<br>CTTTCCGGCTGTTATTGTT<br>GTTATCACTGCTTCTTCTTC<br>CTATCATGTTACCCATCCAA<br>CTATACACCTTAGACTAGT<br>CATCTTATTGATATACATTC<br>CTCCCATCCAACACAACGG | Twist Biosciences |

| | | |
|---|---|---|
| | TATTCTATTTATTTATCCAAT<br>TAGTCATAGTCGTACCACC<br>ATCCAGCACGAAGGTGCCT<br>CTTTAGTAAAGAGTAGAAA<br>GAAGAACCGGATGGGAAAT<br>GTTTTTGTTACAAAAATGAC<br>ACATATTGTAGTGGACAGA<br>AGGAGTGAGACAGACATGA<br>GCAAGCCAATTTGTTTATAA<br>TTTCTCTTCTAGAAAAAAAT<br>ACATTTTTCCATACTTCACT<br>AGTCAAAACCTTTCACCTTT<br>CTAATACATCTCGTAAACCA<br>TAATCTTGATAGTTCTGAGC<br>ATTTCAATACGAAAGCTTCT<br>CACTGTCTAGATCTCTGAC<br>TGAGTGCCCTCATCAAAAG<br>TGCAATCTGTCATCTGTTTC<br>CTCATAATCACGGAGCACT<br>AATTTTTCTCTCTGCGTCTC<br>TATAATCAGATATCTCTCGT<br>CACTAAGAACTTTCCGAAA<br>TGTTTATGCTTCTCATCTGA<br>CCACTTCGGTTCCGCACAA<br>AAAAGTACGGCATTCCAAA<br>AGAAATCTGATCCCCCTCC<br>GTTCATTCGTGGTCCGAGT<br>CGGTGCCACCAGTCGTTGC<br>GCATTGAATATTTGTTTGGT<br>CCGTTCCCCTTCTTCTCCG<br>ACTGCTGACCTCGGGCACT<br>TTGATGACCGGGCCACCAC<br>CTCAGTACCCCTCTATTAC<br>ACCCTCTTTGCCTCCGCGC<br>ATATGACTCCACCCCTTCT<br>CGTGGAAGGCGTGTATCTC<br>CCCTCTTTTCCGCTATTCC<br>CTCGATGGATATATATTCAA<br>ATGTATGTGTGTTCCTGAC<br>GGGAGGGCGTCTCGCTTG<br>AGAGCATCGTCACATCTTT<br>TACAATTTTACTTATGATTT<br>TACTTCATCTTCTTCTTCTT<br>ACTGCGATTTTGATATGCAT<br>TCTTATGTAAACTATTATTA<br>TTCCAGGTTTCCTCACTCTT<br>TTCAAATGAGTAAAGGAGA<br>AGAATTGTTCACTGGAG | |
| *gfp::cki-1* right homology arm | GCGTGATTACAAGGATGAC<br>GATGACAAGAGAATGTCTT | Twist Biosciences |

| | |
|---|---|
| | CTGCTCGTCGTTGCCTTTT<br>CGGTCGTCCGACGCCCGA<br>GCAACGCTCCAGGACTCGA<br>ATTTGGCTTGAAGATGCTG<br>TTAAGCGCATGCGCCAGGA<br>AGAAAGCCAGAAATGGGGA<br>TTCGACTTTGAACTGGAGA<br>CTCCCCTCCCAAGCTCTGC<br>TGGATTCGTTTATGAAGTTA<br>TTCCAGAGAATTGTGTTCC<br>GGAGTTCTACAGGTAATTG<br>AATTTTATAAATTTTTCATA<br>GTTATTTTACTAAACAGTTT<br>CATTTTTCAGAACCAAAGTT<br>CTCACTGTCAGAACCACAT<br>GCTCATCGCTGGACATCAG<br>CTCAACGACTTTGACTCCA<br>TTGAGCTCTCCGAGCACAT<br>CTGATAAGGAGGAGCCCTC<br>GCTGATGGATCCCAACAGC<br>TCGTTCGAAGATGAAGAGG<br>AACCGAAGAAGTGGCAATT<br>CAGAGAGCCACCAACTCCA<br>CGGAAGACCCCAACAAAGC<br>GTCAGCAGAAGATGACCGA<br>CTTCATGGCAGTTTCCCGT<br>AAGAAGAATTCGTTGTCTC<br>CAAACAAGCTGTCTCCGGT<br>GAATGTGATCTTCACTCCA<br>AAATCTCGTCGTCCAACGA<br>TCAGAACTCGATCTTCATG<br>CTCTCCATACTAGAGGTTT<br>CATTTTGACTTTTTTTTGCC<br>CAATTCCACGGGTTGAATC<br>TAATCATTTGATTATCTCCT<br>CGACAGTTTCTGAGTCTCT<br>CTTAATTGTTCAACTAGTCA<br>TGTTTCCACAAATGTTTTAT<br>TGTTTGTTCCAAAAGCCCT<br>GTGATCCATGTTTAGGAAC<br>TCTGTAACTCTTTTTTCCCA<br>TTGCCATTTGTTTTAAACAA<br>CTCAAAGAAAAATAAACCC<br>TTTGAAATTATTTTAAGAAC<br>TGTATTCTGGTGTTTTCTTC<br>AACTTATAAAAAAAAAGACG<br>AATAGAAACTGGCACACGG<br>TGCAGTTCCATTGGTAACT<br>TCAGCAAAGAATATACTGA<br>AATCACGAAAAGTGGTACA |

| | | |
|---|---|---|
| | ATTCCGCGCATAATTTTGAA<br>ACTTCTAACATTCTTCATTA<br>ACTTCAAACTTCAAACATTC<br>TGTAAATGTTGTAAGATCAA<br>ATAAATCTTTCCCGGTTCAC<br>CCACTGCCACCCAAATAGA<br>CATTGCGCGATAACATGGT<br>CATAGCTGTTTCCTGTGTG | |
| *oMS218 (5' HA block)* | ttgtaaaacgacggccagtcgccggca<br>ttatgacaattttctgcgagagtttaaaat<br>attacagatttttttaaattttgaaaaaatatc<br>taatattctcgaaaaaattcgccttggaaa<br>atttcgaaaaaattcattttaaaaatagga<br>aattcaaaattactactttagcattaaaa<br>aaatcataaaaaattctccaaatttttaga<br>agtttccaaaaaaaaaaatcgcaaaaat<br>taaatttgtggttttccaacaataaatgga<br>ccaaaatcaaaaaatttccaccaaaaaa<br>aacataacttctcctcgaggagtacacg<br>agctccgtaaatcgacacagacatttgt<br>gaaaaaaaattacttgaaaatcgtaaaat<br>ttcaacaaaaaaaaattctaatttttttcca<br>gATACTTCCGATTCACCGA<br>CAACAACATTGAAGCAATC<br>AACGAGTTGCTCGATGAAG<br>AGCTCCAAATTACTCAGAA<br>AAAGATTGATGAGCAACGA<br>AACACCCAAATTGCACAAA<br>TGAGCCAtCACCACACACC<br>ACGGCCATCCAAAGCAGCA<br>AGATCTCTCAAATTTCATGG<br>AGCATCGGGAGCCTCAGG<br>AGCATCGATG | Integrated DNA Technologies |
| *DHB-mNG-p2a* | CTTCCATTTCAGGTGTCGT<br>GAACACGCTACCGGTCTCG<br>AGAATTCACCGGATCCATG<br>ACAAATGATGTCACCTGGA<br>GCGAGGCCTCTTCGCCTGA<br>TGAGAGGACACTCACCTTT<br>GCTGAAAGATGGCAATTAT<br>CTTCACCTGATGGAGTAGA<br>TACAGATGATGATTTACCAA<br>AATCGCGAGCATCCAAAAG<br>AACCTGTGGTGTGAATGAT<br>GATGAAAGTCCAAGCAAAA<br>TTTTTATGGTGGGAGAATC<br>TCCACAAGTGTCTTCCAGA<br>CTTCAGAATTTGAGACTGA<br>ATAATTTAATTCCCAGGCAA | Twist Biosciences |

| | | |
|---|---|---|
| | CTTTTCAAGCCCACCGATA<br>ATCAAGAAACTGGTTCCGG<br>GGCCCAGGGCAGCGGCAT<br>GGTGAGCAAGGGCGAGGA<br>GGATAACATGGCCTCTCTC<br>CCAGCGACACATGAGTTAC<br>ACATCTTTGGCTCCATCAA<br>CGGTGTGGACTTTGACATG<br>GTGGGTCAGGGCACCGGC<br>AATCCAAATGATGGTTATG<br>AGGAGTTAAACCTGAAGTC<br>CACCAAGGGTGACCTCCAG<br>TTCTCCCCCTGGATTCTGG<br>TCCCTCATATCGGGTATGG<br>CTTCCATCAGTACCTGCCC<br>TACCCTGACGGGATGTCGC<br>CTTTCCAGGCCGCCATGGT<br>AGATGGCTCCGGATACCAA<br>GTCCATCGCACAATGCAGT<br>TTGAAGATGGTGCCTCCCT<br>TACTGTTAACTACCGCTAC<br>ACCTACGAGGGAAGCCACA<br>TCAAAGGAGAGGCCCAGG<br>TGAAGGGGACTGGTTTCCC<br>TGCTGACGGTCCTGTGATG<br>ACCAACTCGCTGACCGCTG<br>CGGACTGGTGCAGGTCGA<br>AGAAGACTTACCCCAACGA<br>CAAAACCATCATCAGTACC<br>TTTAAGTGGAGTTACACCA<br>CTGGAAATGGCAAGCGCTA<br>CCGGAGCACTGCGCGGAC<br>CACCTACACCTTTGCCAAG<br>CCAATGGCGGCTAACTATC<br>TGAAGAACCAGCCGATGTA<br>CGTGTTCCGTAAGACGGAG<br>CTCAAGCACTCCAAGACCG<br>AGCTCAACTTCAAGGAGTG<br>GCAAAAGGCCTTTACCGAT<br>GTGATGGGCATGGACGAG<br>CTGTACAAGGGCAGCGGC<br>GCCACCAACTTCAGCCTGC<br>TGAAGCAGGCCGGCGACG<br>TGGAGGAGAACCCCGGCC<br>CCATGCCAGAGCCAGCGA<br>AGTCTGCTCCCGCC | |
| *mScarlet-CAAX* | TCTAGAGGCAGCGGCCAG<br>TGCACCAACTACGCCCTGC<br>TGAAGCTGGCCGGCGACG<br>TGGAGAGCAACCCCGGCC | Twist Biosciences |

| | | |
|---|---|---|
| | CCATGGTGAGCAAGGGCG<br>AGGCAGTGATCAAGGAGTT<br>CATGCGGTTCAAGGTGCAC<br>ATGGAGGGCTCCATGAACG<br>GCCACGAGTTCGAGATCGA<br>GGGCGAGGGCGAGGGCC<br>GCCCCTACGAGGGCACCC<br>AGACCGCCAAGCTGAAGGT<br>GACCAAGGGTGGCCCCCT<br>GCCCTTCTCCTGGGACATC<br>CTGTCCCCTCAGTTCATGT<br>ACGGCTCCAGGGCCTTCAC<br>CAAGCACCCCGCCGACATC<br>CCCGACTACTATAAGCAGT<br>CCTTCCCCGAGGGCTTCAA<br>GTGGGAGCGCGTGATGAA<br>CTTCGAGGACGGCGGCGC<br>CGTGACCGTGACCCAGGA<br>CACCTCCCTGGAGGACGG<br>CACCCTGATCTACAAGGTG<br>AAGCTCCGCGGCACCAACT<br>TCCCTCCTGACGGCCCCGT<br>AATGCAGAAGAAGACAATG<br>GGCTGGGAAGCGTCCACC<br>GAGCGGTTGTACCCCGAG<br>GACGGCGTGCTGAAGGGC<br>GACATTAAGATGGCCCTGC<br>GCCTGAAGGACGGCGGCC<br>GCTACCTGGCGGACTTCAA<br>GACCACCTACAAGGCCAAG<br>AAGCCCGTGCAGATGCCC<br>GGCGCCTACAACGTCGAC<br>CGCAAGTTGGACATCACCT<br>CCCACAACGAGGACTACAC<br>CGTGGTGGAACAGTACGAA<br>CGCTCCGAGGGCCGCCAC<br>TCCACCGGCGGCATGGAC<br>GAGCTGTACAAGTCCGGAC<br>TCAGATCTAAGCTGAACCC<br>TCCTGATGAGAGTGGCCCC<br>GGCTGCATGAGCTGCAAGT<br>GTGTGCTCTCCTGACTAGA<br>GTTAACATCGAGGGATCAA<br>GCTTATCGATAATCAACCT<br>CTGGATTACAAAATTTGT | |
| *H2B-mTurqoise2* | ATGCCAGAGCCAGCGAAGT<br>CTGCTCCCGCCCCGAAAAA<br>GGGCTCCAAGAAGGCGGT<br>GACTAAGGCGCAGAAGAAA<br>GGCGGCAAGAAGCGCAAG | Twist Biosciences |

| | |
|---|---|
| | CGCAGCCGCAAGGAGAGC<br>TATTCCATCTATGTGTACAA<br>GGTTCTGAAGCAGGTCCAC<br>CCTGACACCGGCATTTCGT<br>CCAAGGCCATGGGCATCAT<br>GAATTCGTTTGTGAACGAC<br>ATTTTCGAGCGCATCGCAG<br>GTGAGGCTTCCCGCCTGG<br>CGCATTACAACAAGCGCTC<br>GACCATCACCTCCAGGGAG<br>ATCCAGACGGCCGTGCGC<br>CTGCTGCTGCCTGGGGAG<br>TTGGCCAAGCACGCCGTGT<br>CCGAGGGTACTAAGGCCAT<br>CACCAAGTACACCAGCGCT<br>AAGGtTCCACCGGTCGCCA<br>CCATGGTGAGCAAGGGCG<br>AGGAGCTGTTCACCGGGG<br>TGGTGCCCATCCTGGTCGA<br>GCTGGACGGCGACGTAAA<br>CGGCCACAAGTTCAGCGTG<br>TCCGGCGAGGGCGAGGGC<br>GATGCCACCTACGGCAAGC<br>TGACCCTGAAGTTCATCTG<br>CACCACCGGCAAGCTGCC<br>CGTGCCCTGGCCCACCCT<br>CGTGACCACCCTGTCCTGG<br>GGCGTGCAGTGCTTCGCC<br>CGCTACCCCGACCACATGA<br>AGCAGCACGACTTCTTCAA<br>GTCCGCCATGCCCGAAGG<br>CTACGTCCAGGAGCGCAC<br>CATCTTCTTCAAGGACGAC<br>GGCAACTACAAGACCCGC<br>GCCGAGGTGAAGTTCGAG<br>GGCGACACCCTGGTGAAC<br>CGCATCGAGCTGAAGGGC<br>ATCGACTTCAAGGAGGACG<br>GCAACATCCTGGGGCACAA<br>GCTGGAGTACAACTACTTT<br>AGCGACAACGTCTATATCA<br>CCGCCGACAAGCAGAAGA<br>ACGGCATCAAGGCCAACTT<br>CAAGATCCGCCACAACATC<br>GAGGACGGCGGCGTGCAG<br>CTCGCCGACCACTACCAGC<br>AGAACACCCCCATCGGCGA<br>CGGCCCCGTGCTGCTGCC<br>CGACAACCACTACCTGAGC<br>ACCCAGTCCAAGCTGAGCA |

| | | |
|---|---|---|
| | AAGACCCCAACGAGAAGC<br>GCGATCACATGGTCCTGCT<br>GGAGTTCGTGACCGCCGC<br>CGGGATCACTCTCGGCATG<br>GACGAGCTGTACAAGTCTA<br>GAGGCAGCGGCCAGTGCA<br>CCAACTACGCC | |
| *AID-link-miRFP670* | AATACAAGCTACTTGTTCTT<br>TTTGCAGGATCCATCATCC<br>CTTAATTAAGGATAGTGATT<br>ATCGATACATGAAGGAGAA<br>GAGTGCTTGTCCTAAAGAT<br>CCAGCCAAACCTCCGGCCA<br>AGGCACAAGTTGTGGGATG<br>GCCACCGGTGAGATCATAC<br>CGGAAGAACGTGATGGTTT<br>CCTGCCAAAAATCAAGCGG<br>TGGCCCGGAGGCGGCGGC<br>GTTCGTGAAGGTATCAATG<br>GACGGAGCACCGTACTTGA<br>GGAAAATCGATTTGAGGAT<br>GTATAAAGGTGCTAGCGGT<br>GCAGGCGCCATGGTAGCA<br>GGTCATGCCTCTGGCAGCC<br>CCGCATTCGGGACCGCCT<br>CTCATTCGAATTGCGAACA<br>TGAAGAGATCCACCTCGCC<br>GGCTCGATCCAGCCGCAT<br>GGCGCGCTTCTGGTCGTCA<br>GCGAACATGATCATCGCGT<br>CATCCAGGCCAGCGCCAA<br>CGCCGCGGAATTTCTGAAT<br>CTCGGAAGCGTACTCGGC<br>GTTCCGCTCGCCGAGATCG<br>ACGGCGATCTGTTGATCAA<br>GATCCTGCCGCATCTCGAT<br>CCCACCGCCGAAGGCATG<br>CCGGTCGCGGTGCGCTGC<br>CGGATCGGCAATCCCTCTA<br>CGGAGTACTGCGGTCTGAT<br>GCATCGGCCTCCGGAAGG<br>CGGGCTGATCATCGAACTC<br>GAACGTGCCGGCCCGTCG<br>ATCGATCTGTCAGGCACGC<br>TGGCGCCGGCGCTGGAGC<br>GGATCCGCACGGCGGGTT<br>CACTGCGCGCGCTGTGCG<br>ATGACACCGTGCTGCTGTT<br>TCAGCAGTGCACCGGCTAC<br>GACCGGGTGATGGTGTATC | Twist Biosciences |

| | |
|---|---|
| | GTTTCGATGAGCAAGGCCA<br>CGGCCTGGTATTCTCCGAG<br>TGCCATGTGCCTGGGCTCG<br>AATCCTATTTCGGCAACCG<br>CTATCCGTCGTCGACTGTC<br>CCGCAGATGGCGCGGCAG<br>CTGTACGTGCGGCAGCGC<br>GTCCGCGTGCTGGTCGAC<br>GTCACCTATCAGCCGGTGC<br>CGCTGGAGCCGCGGCTGT<br>CGCCGCTGACCGGGCGCG<br>ATCTCGACATGTCGGGCTG<br>CTTCCTGCGCTCGATGTCG<br>CCGTGCCATCTGCAGTTCC<br>TGAAGGACATGGGCGTGC<br>GCGCCACCCTGGCGGTGT<br>CGCTGGTGGTCGGCGGCA<br>AGCTGTGGGGCCTGGTTGT<br>CTGTCACCATTATCTGCCG<br>CGCTTCATCCGTTTCGAGC<br>TGCGGGCGATCTGCAAAC<br>GGCTCGCCGAAAGGATCG<br>CGACGCGGATCACCGCGC<br>TTGAGAGCTAA |

*The ZF degron in CDT-1::ZF::GFP does not cause degradation, because the *zif-1(gk117)* null allele removes the E3 ligase component ZIF-1 that recognizes the ZF tag (Sallee et al., 2018).

#### EXTENDED EXPERIMENTAL PROCEDURES

##### *C. elegans* Transgenic Strain Generation

Transgene insertion was performed via CRISPR/Cas9 genome engineering to generate single copy knock-ins to a known neutral locus on chromosome I or II using a self-excising cassette (SEC)-based method (de la Cova et al., 2017; Dickinson et al., 2015). Homologous repair templates and guide plasmids were graciously provided by Bob Goldstein, targeting the MosSCI integration sites ttTi4348 and ttTi5605 on chromosome I and II, respectively. CRISPR microinjection products were prepared using the PureLink HQ Mini Plasmid DNA Purification Kit from Invitrogen (K210001). An additional wash step was included prior to the final ethanol wash, using 650 µL of 60% 4 M guanidine hydrochloride (Fisher Scientific, BP178-500; pH 6.5, 40% isopropanol) yielding a marked increase in knock-in efficiency. All purified microinjection products were stored at 4˚C.

Injection mixes were freshly made before each round of injection. These mixes contain Cas9-sgRNA plasmids (50 ng/µL), homologous repair templates (50 ng/µL), and a co-injection marker (pCFJ90, 2.5 ng/µL). Injection mixes were injected into the gonads of young adult *C. elegans* N2 hermaphrodites. Successful integrants were identified in the F3 offspring of injected worms (Dickinson et al., 2015). Injected young adult hermaphrodites of the relevant parent strain were then each individually transferred to a fresh OP50 plate and allowed to lay eggs for three days at 25˚C. On day 3, 400 µL of a 5 mg/mL stock of hygromycin B (Millipore, 400052) was added to the plates to a final plate concentration of 0.25 mg/mL. After five days of hygromycin B exposure, surviving dominant *sqt-1* roller (Rol) worms were singled out onto fresh OP50 plates, checked for expression of the desired transgene/genomic edit and the presence of extrachromosomal array markers on a fluorescence dissecting microscope (frame and automation: Zeiss Axio Zoom.V16, light source: Lumencor SOLA light engine). The Rol phenotype was assessed for Mendelian inheritance, and if possible, the genomic edit was homozygosed. Once homozygosed, selectable markers (hygromycin B resistance and dominant *sqt-1* Rol phenotype) were removed from the genome using heat shock-inducible Cre-Lox recombination via either a 3–4 hour heat shock at 34˚C or overnight (8–12 hours) heat shock of large numbers of L1 and L2 stage animals at 26˚C in an air incubator. After two days, wild type worms were singled out one to a plate and progeny assessed for expression and homozygosity of the desired genomic insertion.

##### Zebrafish Transgenic Line Generation

Three transgenic lines were generated, including *Tg(ubb:Lck.mNeonGreen)*$^{sbu107}$, *Tg(hsp70l:DHB.mNeonGreen-p2a-H2B.mScarlet)*$^{sbu108}$, and *Tg(hsp70l:DHB.mScarlet-p2a-H2B.miRFP670)*$^{sbu109}$. These lines were created using the Tol2 transposable element system (Kawakami, 2004). Zebrafish plasmids for generating transgenic lines were created using a *tol2* plasmid vector containing the *hsp70l* promoter based on previous plasmids constructs (Row et al., 2016). For the *hsp70l:DHB.mNeonGreen-p2a-H2B.mScarlet* plasmid, Gibson cloning was used to insert DNA encoding amino acids 994-1087 of human DHB fused to the N-terminus of mNeonGreen, followed by the P2A viral peptide sequence and human H2B with a C-terminal mScarlet fusion. The same method was used to generate *hsp70l:DHB.mScarlet-p2a-H2B.miRFP670*, except mScarlet and miRFP670 were used instead of mNeonGreen and mScarlet, respectively. The *tol2 hsp70l* vector was also used to create the *ubb:Lck.mNeonGreen* plasmid. The *hsp70l* promoter was replaced with the *ubb* promoter (Mosimann et al., 2011), followed by mNeonGreen with an N-terminal membrane targeting sequence from *Mus musculus* LCK (amino acids MGCVCSSNPE). Each plasmid was co-injected with *in vitro* transcribed *tol2* transposase mRNA. One nanoliter of injection mix containing 25 pg/nl of plasmid and 25 pg/nl of tol2 mRNA were injected into wild type zebrafish embryos at the 1-cell stage. Injected embryos were raised to adults and screened for germline transmission.

**Zebrafish Mosaic Analysis**

One nanoliter of injection mix containing 25 pg/nl of HS-PCNA-GFP and 25 pg/nl of tol2 mRNA were injected into *Tg(hsp70l:DHB.mScarlet-p2a-H2B.miRFP670)*$^{sbu109}$ zebrafish embryos at the 1-cell stage.

**Molecular Biology**

Synthetic DNAs were ordered as gBlocks from Integrated DNA Technologies (IDT) or gene fragments from Twist BioScience (see Key Resources Table). The nucleotide sequence of DHB (index 1.0) was codon optimized for *C. elegans* somatic expression and the P2A sequence used in pWZ193 (index 0.2; see KRT) de-optimized to increase the efficiency of ribosome stalling (Lo et al., 2019; Redemann et al., 2011). The *C. elegans* *rps-0* and *rps-27* promoters and the *pcn-1* promoter and coding sequence were all amplified from N2 genomic DNA. Sequences of all primers and synthetic DNAs are provided in the KRT. Synthetic gene fragments and amplified DNAs were cloned via Gibson Assembly (Barnes, 1994; Gibson et al., 2010; Gibson et al., 2009) or NEBuilder HiFi into target plasmids.

Constructs used for zebrafish transgenes were made from PCR products amplified from synthetic Twist BioSience gene fragment sequences followed by NEBuilder HiFi cloning. Human DHB and H2B sequences were used for making the DHB transgenes, and the human membrane targeting Lck sequence was used for the *ubb:Lck.mNeonGreen* transgene. All primers and synthetic gene fragment sequences are available in the KRT.

**Microinjection Setup**

Microinjections for *C. elegans* transgenesis were performed on an injection setup combining a Zeiss Axio Observer A1 inverted compound frame, EC Plan-Neofluar 40x/0.75 NA DIC objective and floating stage, with a Narashige manual micromanipulator and a picoliter injection system from Warner for fine control of delivered volume. Microinjection needles (Sutter) were pulled on a Sutter P-97 reconditioned and calibrated by Sutter.

Zebrafish microinjections were performed on either a Leica S6e or a Zeiss Stemi 508 dissecting microscope using a Narishige manual micromanipulator and a Warner picoliter injecting system. Glass needles were pulled on a P-1000 puller from Sutter Instruments.

***C. elegans* RNAi Perturbations**

RNAi was delivered by feeding *E. coli* strain HT115(DE3) expressing double-stranded RNA (dsRNA) to synchronized L1 stage strains. Transcription of dsRNA was induced with 1 mM isopropyl b-D-1-thiogalactopyranoside (Thermo Scientific, R0393) in bacterial cultures for one hour at 37˚C. After an hour, cultures were plated on NGM plates topically treated with 2.5 µl each of 30 mg/mL carbenicillin (Alfa Aesar, J61949) and 10 µl of 1 M IPTG (Kelley et al. 2019). The RNAi vector targeting *cdk-1* was obtained from the Ahringer RNAi library (Kamath et al., 2003). The empty vector L4440 was used as a negative control. RNAi vectors were verified by Sanger sequencing.

##### *C. elegans* CKI-1 Experiments

For heat shock CKI-1 experiments, the following strains were used DQM406 (*hsp*>CKI-1::BFP; *rps-0*>DHB::mKate2) and DQM394 (*rps-0*>DHB::mKate2). Synchronized L1 animals were plated on OP50 and allowed to develop to late L2/early L3. Plates were then placed at 30˚C in an air incubator for 3 hours. Animals were then placed at 20˚C and allowed to recover from heat shock for 20-40 min before being mounted for static imaging.

For assessing endogenous CKI-1 levels in Figure 4, strain DQM586 (GFP::CKI-1; *rps-27*>DHB::2xmKate2) was utilized. Briefly, L1 animals were synchronized via sodium hypochlorite treatment and plated on OP50 at 25˚C and analyzed at the P6.p 2-cell, 4-cell, and 8-cell stages. DQM586 was superficially wild type, but several phenotypes, revealed by confocal microscopy and/or analyzed in this study (e.g., the presence of larger somatic cells than normal in the L3 and L4 stages, including the anchor cell), led to the conclusion that the N-terminal GFP fusion (which lacks a flexible linker) resulted in animals displaying a gain-of-function effect of GFP::CKI-1. Early terminal differentiation in the VPCs was determined by lineage analysis of each image and comparing the size of individual VPCs in GFP::CKI-1 animals to wild type. A VPC was considered to have undergone early terminal differentiation if it failed to divide (larger nucleus than normal) and showed strong nuclear localization of DHB::2xmKate2 consistent with a $CDK^{low}$ state.

##### Zebrafish Drug Perturbations

Palbociclib (PD-0332991), a selective inhibitor of CDK4/6, was purchased from MedChemExpress (HY-A0065). A 5 mM stock solution in embryo media was prepared and stored at -80˚C for up to six months. Prior to each experiment, palbociclib was thawed and diluted in embryo media to a final concentration of 50 µM. Control experiments were performed by treating zebrafish embryos with embryo media only. Embryos were placed in palbociclib at 16 somites for five hours at 22 ˚C.

##### Microscopes for Live-Cell Imaging

All live-cell imaging of *C. elegans* and zebrafish, unless indicated otherwise, was performed on a custom-assembled spinning disk confocal microscope consisting of a Zeiss Axio Imager A2 frame, a Borealis modified Yokogawa CSU10 spinning disc, an ASI 150 micron piezo stage controlled by a MS2000, an ASI filter wheel and a Hamamatsu ImagEM X2 EM-CCD camera. The imaging objective used for *C. elegans* imaging was a Plan Apochromat 100x/1.4 NA DIC objective (Carl Zeiss). For zebrafish imaging, a Plan Apochromat 63x/1.0 NA water dipping objective (Carl Zeiss) was used. Animals in Figure 1 were imaged on a separate custom-assembled spinning disk confocal microscope consisting of an automated Zeiss frame, a Yokogawa CSU10 spinning disc, a Ludl stage controlled by a Ludl MAC6000 and an ASI filter turret attached to a Photometrics Prime 95B camera. The imaging objective used was a Plan Apochromat 63x/1.4 NA DIC objective (Carl Zeiss). For both aforementioned microscopes, laser illumination was provided by a six-line, 405/442/488/514/561/640 nm Vortran laser merge driven by a custom Measurement Computing Microcontroller integrated by Nobska Imaging, Inc. Both microscopes were controlled with Metamorph software (version: 7.10.2.240) and laser power levels were set with Vortran's Stradus VersaLase 8 software. In Figure 1 and S1, live imaging of *C. elegans* embryos was performed on a Nikon Ti-E inverted microscope using a Plan Apochromat 60x/1.4 NA oil immersion objective and controlled by Nikon's NIS-Elements software (version: 4.30). Images were acquired with an Andor Ixon Ultra back thinned EM-CCD camera using 488 nm or 561 nm imaging lasers and a Yokogawa X1 confocal spinning disk head equipped with a 1.5 Å magnifying lens. For time-lapse imaging of the *C. elegans* germline and embryos in Figure S1 and Movie S1, recordings were acquired using a Yokogawa CSUW1 SoRa spinning disk confocal in SoRa disk mode with 1.0x relay lens, a 60x/1.27 NA water immersion objective and a Prime 95B sCMOS camera mounted on a Nikon Ti-2 stand. Nikon's NIS-Elements software (version: 4.3) was used for image acquisition.

##### ***C. elegans*** **Imaging Conditions**

For static imaging experiments, worms were anesthetized by mounting on a 7.5% noble agar pad containing sodium azide (Sigma-Aldrich, S2002) (Martinez and Matus, 2020; Matus et al., 2015). Time-lapse imaging of *C. elegans* was performed using a modified version of a previously published protocol (Kelley et al., 2017). We substituted in a 24 mm square coverslip #1.5 (Fisher Scientific, 12-541-B) and divided the imaging agar pad into two asymmetric smaller portions (each 2-3 mm squares), filling the void space under the coverslip with 5 mM levamisole in M9 buffer or M9 buffer alone. These modifications allowed for much longer imaging durations and substantially reduced sample Z-drift over the course of the imaging session on both upright and inverted microscope systems.

Anesthesia was performed in a spot dish in ~50 µl of a 0.1% tricaine (Sigma-Aldrich, E10521)/0.01% levamisole hydrochloride (Sigma-Aldrich, L9756) anesthetic (Kirby et al., 1990; Maddox and Maddox, 2012; Wong et al., 2011). For some experiments, this tricaine-levamisole solution was substituted for 5 mM levamisole in M9 buffer. When levamisole was used alone, to maintain animals in an anesthetized state for long-duration time-lapse imaging, imaging chambers were flooded with 5 mM levamisole in M9 instead of M9.

Embryos for imaging (Figure S1A, S1B, Movie S1) were collected by dissection from gravid hermaphrodites and incubated for 4–4.5 hours in M9 at room temperature (Figure S1) or imaged immediately (Figure S1B, Movie S1). For live imaging, images were taken at a sampling rate of 0.5 µm. For time-lapse, Z-stacks were collected every four (Figure S1) or three min (Figure S1B, Movie S1). For time-lapse of the germline (Figure S1D, Movie S1), young adult worms were lightly immobilized using 0.1 mM levamisole in M9 buffer and mounted on 5% agarose pads.

##### Zebrafish Imaging Conditions

Zebrafish were mounted in a 35 mm glass bottom dish with uncoated #1.5 coverslip and 20 mm glass diameter (MatTek). A thin layer of 1% agarose dissolved in embryo media (Westerfield, 2007), was added to the dish covering the glass bottom. Once solidified, a P10 pipette tip was used to punch holes in the agarose. Embryos were added to 1% low melting point agarose dissolved in embryo media containing 1x tricaine (24x stock 0.4g/l; Pentair, TRS1), and then one embryo was added to each of the punched holes. Embryos were manipulated gently with an eyelash while the agarose solidified to ensure proper orientation. For 72 hpf embryos, animals were anesthetized in 1x tricaine prior to mounting in 1% low melt agarose with 1x tricaine. In all cases imaging dishes were filled with embryo media containing 1x tricaine.

##### Image Processing

Hand quantification of images was performed in Fiji (version: 2.0.0-rc-69/1.52p) (Schindelin et al., 2012). Due to the high level of amplifier noise in EM-CCD images, and to remove any remaining out-of-focus fluorescence in these confocal micrographs, a rolling ball background subtraction was used (size = 50) (Sternberg and Corporation, 1983). After a recording was qualified for inclusion, ratiometric measurements were obtained.

First, the Z plane containing the center of the cell of interest was located. Using the freehand tool, a conservative toroid was drawn around the nucleus and excluding the nucleolus if present, which does not localize the CDK sensor. The fluorescent histone and corresponding DIC and DHB images were used to assess the accuracy of this toroid. A measurement of mean gray value was obtained. Then, a region of perinuclear cytoplasm was chosen, avoiding pixels belonging to the cytoplasm of neighboring cells. The mean grey value of the cytoplasmic patch was then measured. These values were recorded and a cytoplasmic:nuclear ratio was calculated. If there were multiple cells of interest in the image, the procedure was repeated for each cell. For time-lapse recordings, this procedure was repeated at each time point.

##### Statistical Analyses

To evaluate the predictability of CDK activity (readout as the ratio of cytoplasmic-to-nuclear intensity of DHB) on pre-terminal versus terminal cell fate in different cell cycle phases, we created a receiver operating characteristic (ROC) curve for CDK activity at each time point relative to anaphase. Using the *perfcurve* function in MATLAB, we calculated the area under the curves (AUC) as the indicator of predictability. We then built a classifier to predict pre-terminal vs. terminal cell fates based on CDK activity after anaphase. For each time point, we chose the CDK activity threshold for classification that maximizes the geometric mean of specificity (1 – false positive rate) and sensitivity (true positive rate). We tested the classifier in a second dataset, the stochastic division of the vulval D cell (see **Figure 5**). To predict the cell fate of each trace, we made independent classifications on each relevant time point based on CDK activity and use the majority class of all relevant time points as the classification for the trace. For traces recorded beyond 60 min after anaphase, we used all time points after 60 min post-anaphase, since these time points allow near-perfect prediction (AUC>0.9). For traces recorded beyond 20 min but within 60 min after anaphase, we used their last three time points, since these time points show good and increasing prediction power with AUC>0.8.

Bootstrapping was performed in MATLAB R2019A. The code used is available at GitHub (https://github.com/abraham-kohrman/matus-dhb-stats). Custom code for statistical testing may not be compatible with MATLAB releases older than R2019A and may require the use of MATLAB Toolboxes. Briefly, when single timepoint samples did not exhibit normal distributions, empirical statistics were calculated. For single timepoint experiments, a bootstrapped distribution of the difference between mean groups was calculated for each comparison (Equation 1).

Equation 1: $|\bar{x}_1 - \bar{x}_2|$

$10^8$ statistical simulations were performed by random sampling without replacement in MATLAB. A *p*-value was calculated by determining the proportion of simulated differences with values greater than the true difference.

For comparisons of time course data, a mean trend line was calculated for each dataset to be compared. The area between the mean trend lines was calculated. In MATLAB, this was performed as the sum of the absolute value of the difference at each time point. Where $x_1$ corresponds to the first trend line and $x_2$ corresponds to the second trend line.

Equation 2: $\|x_1 - x_2\|_1$

Statistical simulations were performed by random partitioning of the data without replacement into two groups with the same sizes as the original groups. Mean trend lines were then calculated for these randomly assigned groups, and as before the statistic was calculated. $10^8$ simulated replicates were performed to estimate the distribution of the difference statistic. In a manner analogous to bootstrapping, *p*-value was calculated by determining the proportion of simulations with more extreme statistical values than the observed statistic. See Figure S6 for a detailed schematic of the procedure.

**Reporting of Statistical Results**

The α value for this study was nominally 0.05, however exact *p*-values and *n* (number of cells) are reported in all cases. When no simulation produced a more extreme result than the true data configuration, *p*-values are reported as $p<1\text{x}10^{-7}$, rather as the true probability value is so small, as to be outside the range of accurately calculable probability values. For every comparison performed, plots of distributions of empirically calculated statistics are available upon request.

To interpret *p*-values as presented, it is important to note our null hypothesis which can be formulated as: The categorization (e.g. into C lineage vs. D lineage cells or treated vs untreated cells) is not better than random. In short, the *p*-values we have correspond to the probability that the difference between the mean or mean trend lines arose by chance. Another formulation would be the odds that the categorization of the data is meaningless.

Throughout the study, an α value of 0.05 is used for significance. A *p*-value of 0.05 corresponds to the statement that 95% of random reassortments of the data yielded a difference between the means/mean trend lines less extreme than the true, observed difference.

In the course of data collection for this manuscript, many animals were recorded that were not included in this manuscript. In order to be considered for analysis, recordings had to satisfy the following criteria: (1) a cell of interest had to have been present in the recording, (2) the cell of interest must have exhibited at least one anaphase during the recording, and (3) the animal must have appeared phenotypically normal at the beginning and end of the recording. Additional criteria for exclusion were the presence of a stalled metaphase plate at any point in the movie or unexpected developmental arrest.

**Computational Resources**

For data analysis, two workstation computers were used. Both systems boot into Windows 10 (Microsoft) off a 1 TB M.2 drive (Samsung 970 EVO Plus). The first system consists of an I9-9900X processor (Intel), a GeForce GTX 1070 Ti GPU (Nvidia) and 128 GB of DDR4 RAM (Corsair). The second system has an I9-9900K processor (Intel), a GeForce RTX 2070 GPU (Nvidia) and 64 GB of DDR4 RAM (G.Skill Ripjaws). Data were stored on a 4 TB RAID0 array consisting of two 2 TB drives (Samsung) and a 2 TB RAID0 array consisting of two 1 TB Drives (Samsung), respectively. System integration, support and maintenance performed by Nobska Imaging, Inc.

**Generation of Figures and Supplemental Movies**

Data for figures were plotted in GraphPad Prism (version: 8.1.2). Micrographs in all figures were reviewed and selected in Fiji. Figure micrographs were contrast and brightness adjusted for ease of display in Adobe Photoshop CC (version: 20.0.6) or Fiji. Figures were assembled in Adobe Illustrator CC (version: 23.0.26). Supplemental movies were selected in Fiji and clipped to the desired length. The plane of interest was selected, and a time-lapse montage of channels was created. Time-lapse movies were rotated to standard orientation, cropped to the relevant region and timestamps and scale bars annotations were added. Brightness and contrast were adjusted for ease of viewing. Movies showing more than one channel were assembled using the multi-stack montage plugin (https://github.com/BIOP/ijp-multi-stack-montage).

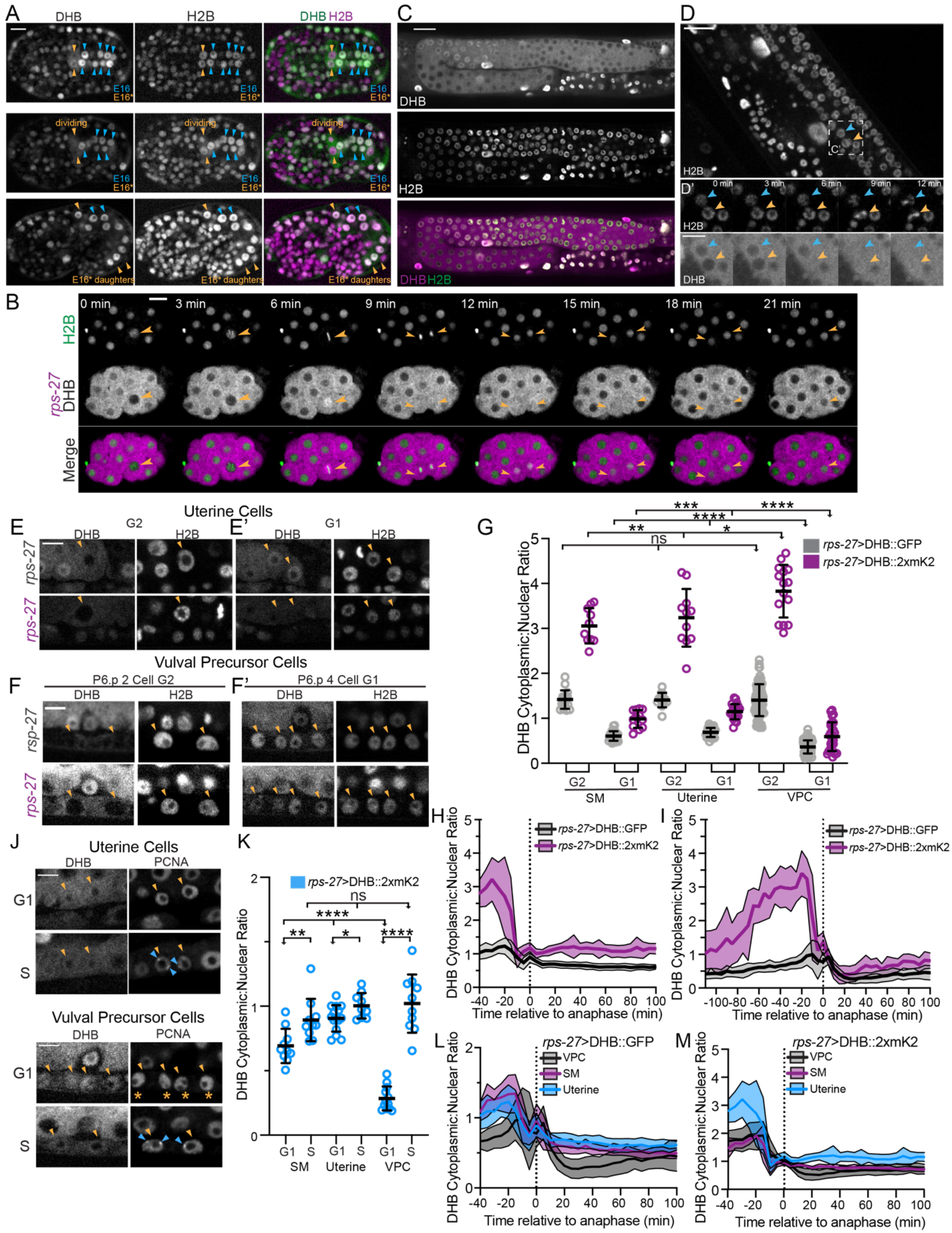

A
DHB
H2B
DHB H2B
E16
E16*
dividing
E16* daughters
C
DHB
H2B
DHB H2B
D
H2B
C'
D'
0 min
3 min
6 min
9 min
12 min
DHB
B
0 min
3 min
6 min
9 min
12 min
15 min
18 min
21 min
H2B
rps-27 DHB
Merge
Uterine Cells
E
G2
E'
G1
rps-27
Vulval Precursor Cells
F
P6.p 2 Cell G2
F'
P6.p 4 Cell G1
rsp-27
G
rps-27>DHB::GFP
rps-27>DHB::2xmK2
DHB Cytoplasmic:Nuclear Ratio
ns
G2
G1
SM
Uterine
VPC
H
I
Time relative to anaphase (min)
J
Uterine Cells
PCNA
S
K
rps-27>DHB::2xmK2
L
rps-27>DHB::GFP
VPC
SM
Uterine
M

**Figure S1. Visualization of cell cycle state live in embryonic and post-embryonic tissues.** *Related to Figure 1*. (A) *C. elegans* embryos expressing the CDK sensor (*rps-27*>DHB::GFP::P2A::H2B::2xmKate2) at the intestinal E16/bean stage through the intestinal E20/1.5-fold stage. Twelve E16 cells arrest (blue arrows) and the remaining four E16* star cells (orange arrows) re-enter the cell cycle and go on to divide, giving rise to four anterior (one shown) and four posterior (two shown) daughters. All 20 intestinal cells remain arrested for the rest of embryonic development ($n \geq 14$ embryos examined). Scale bar = 5 µm. (B) Representative fluorescence overlay (bottom), H2B (top), and DHB::2xmKate2 (middle) time series images during embryo cell divisions (see Movie S1). Orange arrowheads follow the division of a single blastomere. Scale bar, 10 µm. (C) Representative micrograph of young adult germline expressing DHB::2xmKate2 (top), H2B (middle), and overlay (bottom). Scale bar = 10 µm. (D) Representative micrograph and time series insets of H2B and DHB localization in a 12 min window (D'), showing strong nuclear exclusion of DHB::2xmKate2 prior to mitosis in two representative germline nuclei (orange and cyan arrows; see Movie S1). Scale bar = 10 µm (D) and 5 µm (D'). (E, F) Representative images of sensor expression in uterine cells (E) and vulval precursor cells (VPCs; F) at peak G2 and 20 min after anaphase during G1 (E' and F') in DHB::GFP (grey) and DHB::2xmKate2 (magenta). Orange arrowheads denote cells of interest. Scale bar = 5 µm. (G) Dot plot depicting dynamic ranges of the two CDK sensor variants, measured by the cytoplasmic:nuclear ratio of DHB mean fluorescent intensity, at the peak of G2 and G1 in the SMs, uterine cells, and VPCs ($n \geq 10$ cells for each lineage). Scale bar = 5 µm. (H, I) Plot of DHB cytoplasmic:nuclear ratio in uterine cells (H) and VPCs (I) during one round of cell division, measured every 5 min ($n \geq 4$ cells per strain). (J) Representative micrographs of DHB::2xmKate2 and PCNA (*pcn-1*>PCN-1::GFP) in uterine cells (top) and VPCs (bottom) in G1 and S phase (blue arrowheads denote PCN-1::GFP puncta). (K) Dot plot depicting DHB::2xmKate2 ratios during G1 and S phase ($n \geq 9$ cells per phase). (L, M) Plot of DHB cytoplasmic:nuclear ratios in DHB::GFP (L) and DHB::2xmKate2 (M) compared between post-embryonic lineages. Dotted line indicates time of anaphase. Error bars and shaded error bands depict mean±SD. In all supplemental figures: ns, not significant, *$p \leq 0.05$, **$p \leq 0.01$, ***$p \leq 0.001$, ****$p \leq 0.0001$. Significance determined by statistical simulations; *p*-values in Table S1.

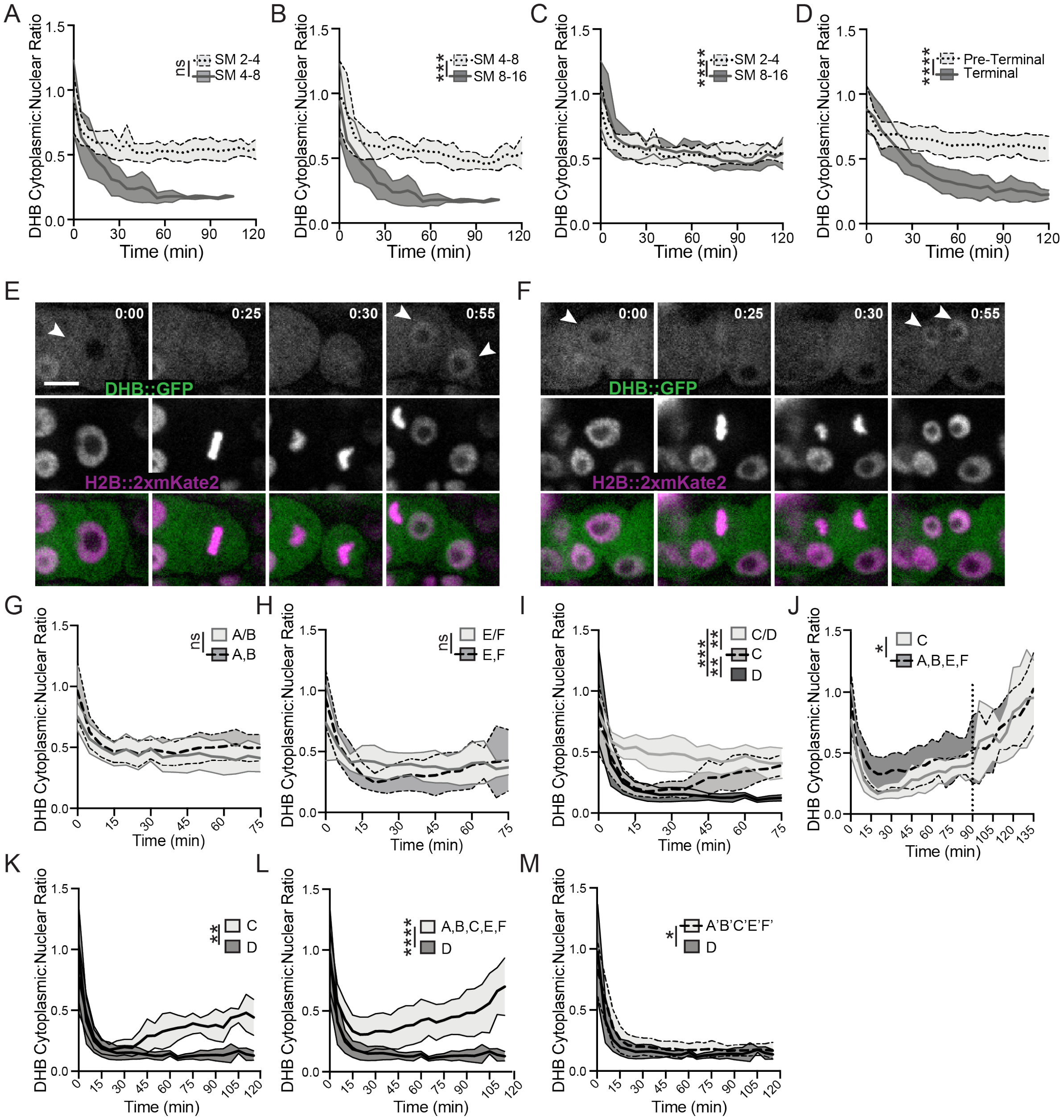

**Figure S2. CDK sensor can accurately detect differences in proliferation potential of postembryonic blast lineages.** *Related to Figure 2, 3, Movie S3-S5*. (A) Cells exit the first and second pre-terminal SM divisions at comparable CDK$^{inc}$ states. (B, C) Cells exit the third and terminal SM division in a CDK$^{low}$ state that is distinct from that of the earlier divisions. (D) DHB ratios in SS/VU cells, comparing pre-terminal cells, which exit in a CDK$^{inc}$ state, and terminal cells, which exit in a CDK$^{low}$ state ($n$≥10 each). (E, F) Stills of time-lapse movies (see Movie S3) showing CDK sensor localization in cycling SS (E) and terminally differentiated sheath cells (F). Scale bar = 5 µm. (G-M) Comparison of CDK activity in pre-terminal and terminal VPC divisions. (G) Comparison of pre-terminal AB ($n$=18) versus A and B cells ($n$=9 each). (H) Comparison of pre-terminal E/F ($n$=13) and E and F cells ($n$=33 cells). (I) Comparison of pre-terminal CD ($n$=10) versus C and D cells ($n$=10 each). (J) Comparison of pre-terminal C ($n$=10) vs. ABEF ($n$=51 cells). (K) Comparison of C vs. D cells ($n$=10 each). (L) Comparison of D ($n$=10) vs. ABCEF cells ($n$=61). (M) Comparison of terminal D cells ($n$=10) vs. A'B'C'E'F' ($n$=94). Time stamp hr:min, scale bar = 10 µm. Shaded error bands depict mean±SD.

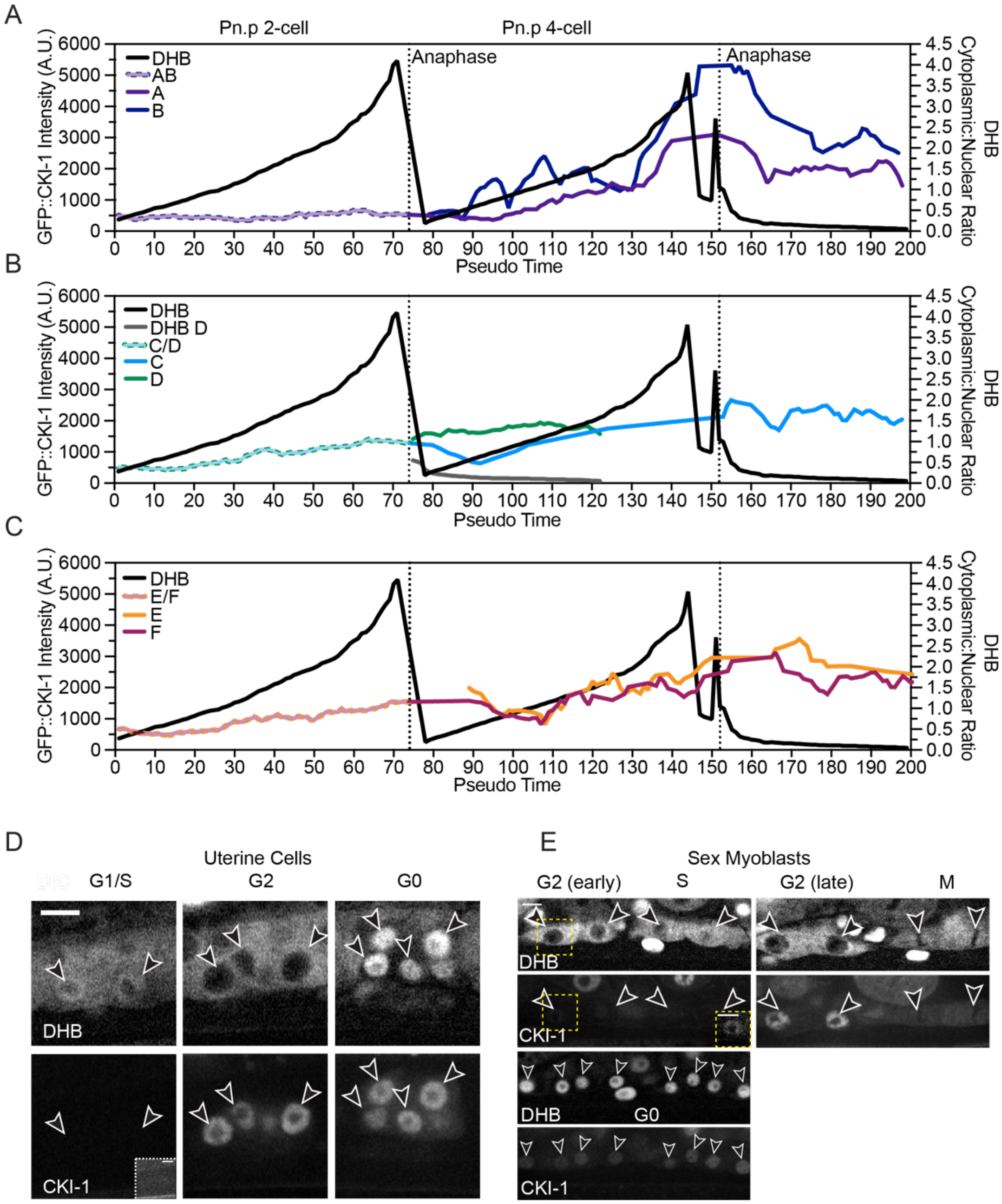

**Figure S3. GFP::CKI-1 levels are predictive of future cell behavior.** *Related to Figure 4*. (A-C) Pseudo-time plots comparing levels of GFP::CKI-1 and CDK sensor ratio in vulval AB (A), CD (B) and EF (C) lineages. (D) Representative single plane confocal micrographs show DHB (top) and GFP::CKI (bottom) localization in uterine cells in pre-terminal (left, middle) and terminal cells (G0, right). (E) Representative single plane confocal micrographs show DHB (top) and GFP::CKI (bottom) localization in SM cells lineages in pre-terminal (top) and terminal cells (G0, bottom). CKI-1 levels increase later in G2 (right). Inset boxes shown for CKI-1 images are contrast enhanced. Scale bar = 5 μm.

A

| Genotype | Pn.p lineages | | | | |
|---|---|---|---|---|---|
| | **P5.p** | **P6.p** | **P7.p** | **°C$^{i}$** | **°C$^{f}$** |
| wild type | LLTU | TTTT | UTLL | 20 | 20 |
| *rps-27>DHB::GFP::P2A::H2B::2xmKate2* | LLTL | TTTT | UTLL | 25 | 25 |
| *rps-27>DHB::GFP::P2A::H2B::2xmKate2* | LLTT | TTTT | ULLL | 25 | 25 |
| *rps-27>DHB::GFP::P2A::H2B::2xmKate2* | LLTU | TTTT | TTLL | 25 | 25 |
| *rps-27>DHB::GFP::P2A::H2B::2xmKate2* | LLLU | TTTT | TLLL | 25 | 25 |
| *rps-27>DHB::GFP::P2A::H2B::2xmKate2* | LLTL | TTTT | UTLL | 25 | 25 |
| *rps-27>DHB::GFP::P2A::H2B::2xmKate2* | LLTT | TTTT | ULLL | 25 | 25 |
| *rps-27>DHB::GFP::P2A::H2B::2xmKate2* | LLTL | TTTT | UTLL | 25 | 25 |
| *rps-27>DHB::GFP::P2A::H2B::2xmKate2* | LLTT | TTTT | UTLL | 25 | 25 |
| *rps-27>DHB::GFP::P2A::H2B::2xmKate2* | LLTU | TTTT | TTLL | 25 | 25 |
| *rps-27>DHB::GFP::P2A::H2B::2xmKate2* | LL†UU | TTTT | TTLL | 25 | 25 |

L: Longitudinal; T: Transverse; U: Undivided, $G_0$ Arrest; †U; Undivided, $G_2$

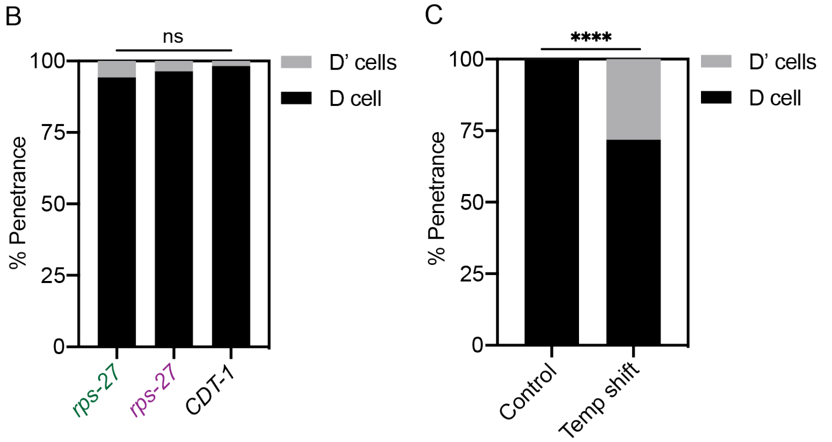

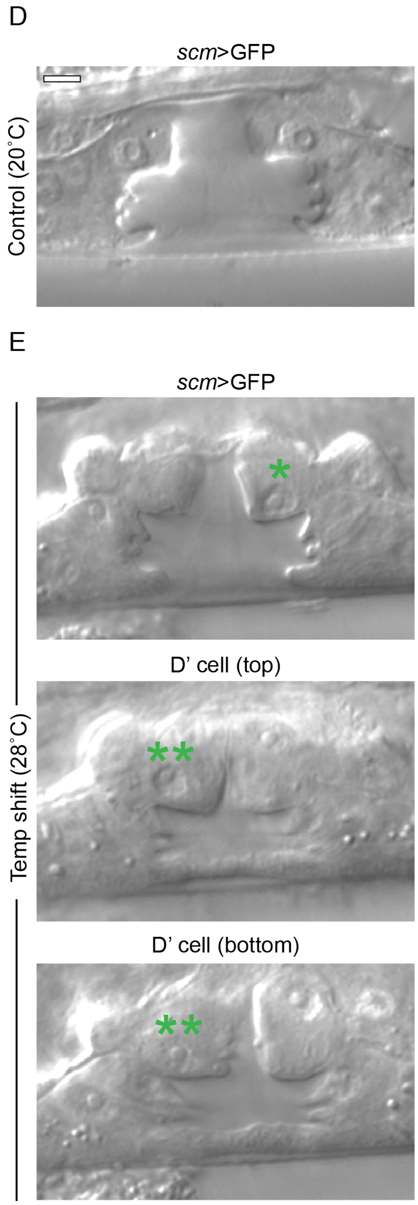

**Figure S4. The vulval D cell divides stochastically.** *Related to Figure 5*. (A) Pn.p lineages for respective genotypes listed. Letters indicate whether and in which orientation the four granddaughters further divided, following the nomenclature of (Sternberg and Horvitz, 1986). Purple text indicates deviation from wild type condition. (B, C) Bar graphs displaying penetrance of D' division in animals grown at 25˚C expressing different variants of the CDK sensor or endogenously tagged CDT-1::ZF::GFP or from temperature shifts from 20-28˚C (C). (D, E) Representative DIC micrographs of 20˚C control (D) and 28˚C experimental (E) conditions of L4 stage vulva with an extra D cell (D') division (**) as compared to wild type (*). Scale bar = 5 μm.

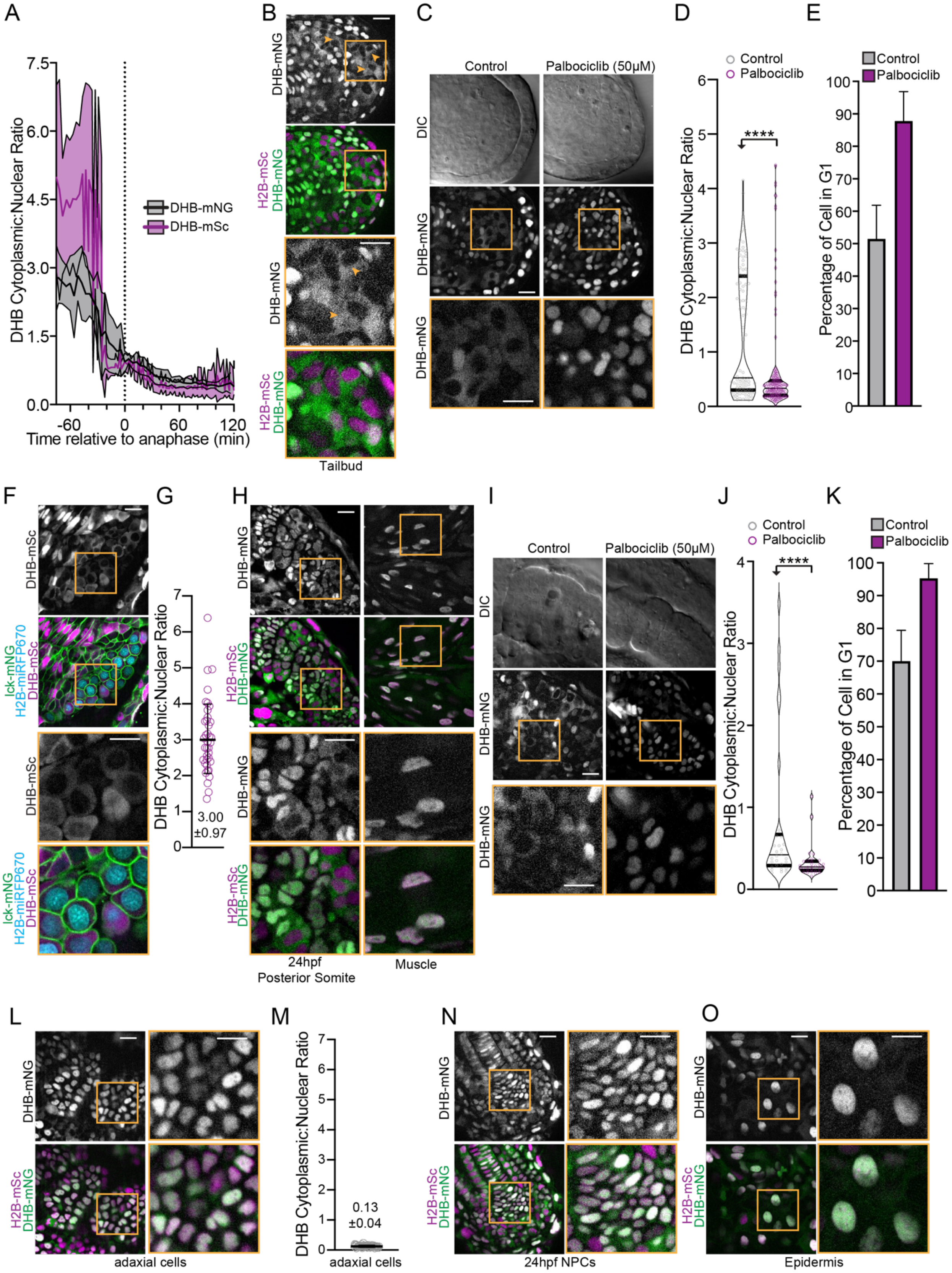

A
B
C
D
E
DHB Cytoplasmic:Nuclear Ratio
Time relative to anaphase (min)
DHB-mNG
DHB-mSc
H2B-mSc
Tailbud
Control
Palbociclib (50μM)
DIC
Palbociclib
Percentage of Cell in G1
F
G
H
I
J
K
lck-mNG
H2B-miRFP670
3.00
±0.97
24hpf
Posterior Somite
Muscle
L
M
N
O
adaxial cells
0.13
±0.04
24hpf NPCs
Epidermis

**Figure S5. CDK sensor expression in zebrafish and CDK4/6 inhibition in developing zebrafish increase percentage of cells in G1.** *Related to Figure 6*. (A) DHB ratio plot from time lapse of DHB-mNG (grey) and DHB-mSc (magenta) of peak G2 through anaphase and G1 ($n$>10 examined for each). Dotted line indicates time of anaphase. (B) Representative micrographs of zebrafish tailbud localization of DHB-mNG. (C) Representative images of the tailbud of control or 50 μm palbociclib treated embryos ($n$≥3 embryos). (D) Quantification of DHB in the tailbud (posterior wall and notochord cells excluded) of control or 50 μm palbociclib treated embryos at 20-22 somite stage. (E) Percentage of cell in G1 in the tailbud (posterior wall and notochord cells excluded) of control or 50 μm palbociclib treated embryos. (F) DHB-mSc expression in primitive red blood cells in the intermediate cell mass at 24 hpf. Inset shows exclusion of DHB from the nucleus indicating G2. (G) Quantification of DHB ratio confirms cells are in G2 with a mean ratio of 3.00. Line and error bars depict mean±SD. (H) Representative micrographs of DHB-mNG in the posterior forming somite (left) and terminally differentiated muscle (right). (I) Representative images of the tailbud of control or 50 μm palbociclib treated embryos ($n$≥3 embryos). (J) Quantification of DHB in the posterior forming somite of control or 50 μm palbociclib treated embryos at 20-22 somite stage. (K) Percentage of cell in G1 in the posterior forming somite of control or 50 μm palbociclib treated embryos. (L) Representative image of the adaxial cells at the 22-somite stage in the most recently formed (posterior) somites. (M) Quantification of DHB ratio confirms cells are in G1 with a mean ratio of 0.13. Line and error bars depict mean±SD. (N, O) Representative micrographs of DHB-mNG localization in notochord progenitor cells (NPCs, N) and epidermis (O). Orange boxes, insets. Scale bar = 20 μm.

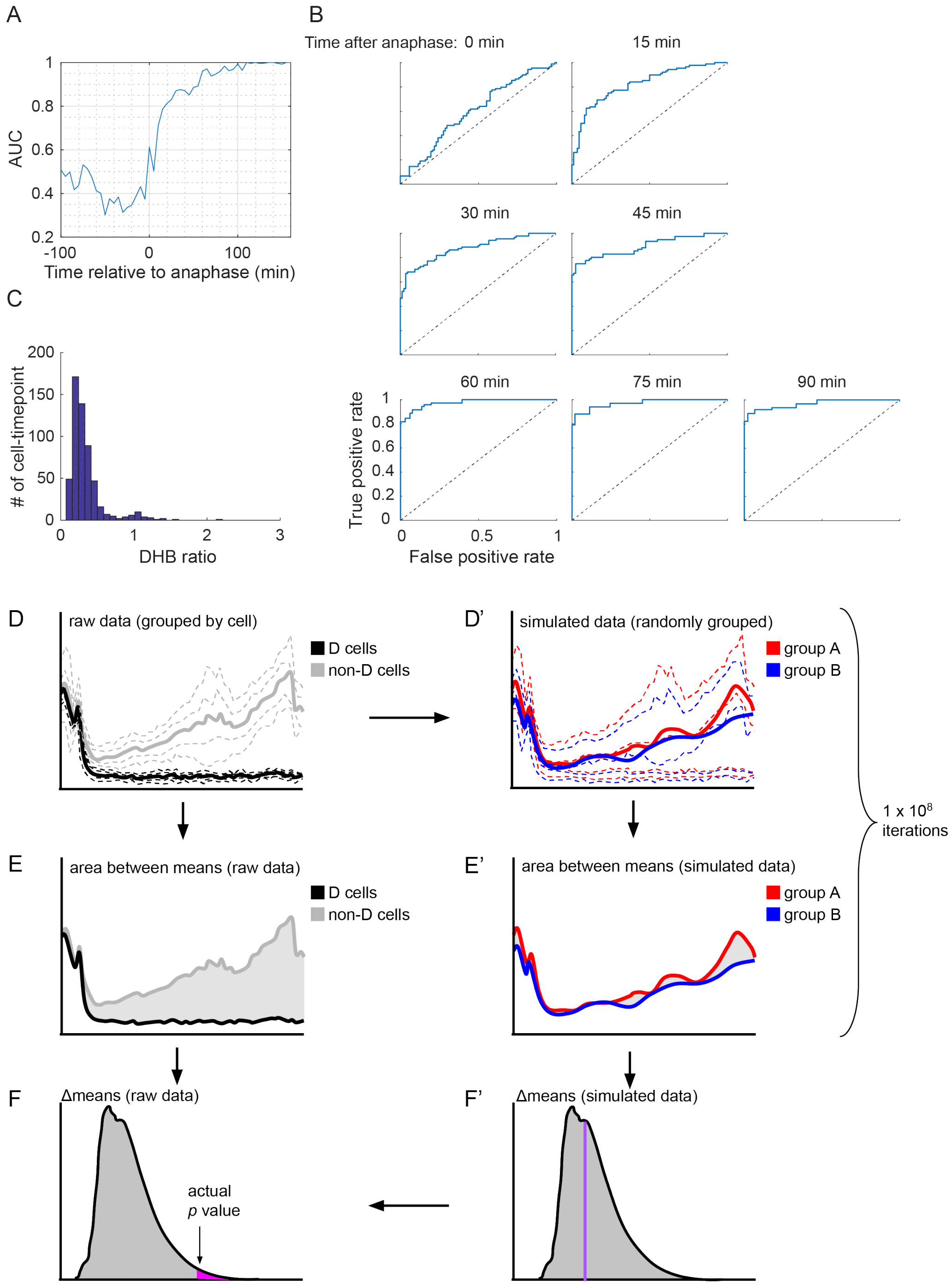

A
B
Time after anaphase: 0 min
15 min
30 min
45 min
60 min
75 min
90 min
AUC
Time relative to anaphase (min)
C
### of cell-timepoint
DHB ratio
True positive rate
False positive rate
D
raw data (grouped by cell)
D cells
non-D cells
D'
simulated data (randomly grouped)
group A
group B
1 x 10^8
iterations
E
area between means (raw data)
E'
area between means (simulated data)
F
Δmeans (raw data)
actual
p value
F'
Δmeans (simulated data)

**Figure S6. Statistical evaluation of the predictive model for future cell behavior in *C. elegans* and zebrafish and a schematic describing the method used for statistical simulations.** *Related to Figure 7*. (A) Predictability of CDK activity on cell fate over time since anaphase. To evaluate the predictability of CDK activity (readout as the ratio of cytoplasmic-to-nuclear intensity of the CDK sensor) on pre-terminal vs. terminal cell fate in different cell cycle phases in *C. elegans*, we created a Receiver Operating Characteristic (ROC) curve for CDK activity at each time point relative to anaphase. Using the *perfcurve* function in MATLAB, we calculated the area under the curve (AUC) as the indicator of predictability. (B) Example ROCs that are used to calculate AUC in (A). (C) A histogram of all data points aggregated ≥50 min after anaphase shows that DHB ratios follow a bimodal distribution, suggesting the presence of two populations. (D) Raw data for each cell type/lineage are grouped together and mean time courses generated. (E) The area between these mean trendlines is then calculated. (D') The assignment between experimental condition groups is randomized, mean time courses are calculated, and (E') the area between the curves is calculated and recorded (purple line in F'). The randomization procedure, (D') and (E'), is repeated 100,000,000 times generating a simulated distribution of the difference statistic: (F'). The true value of the difference statistic is compared to the distribution in (F') generating a *p*-value (F) corresponding to the proportion of values more extreme than the observed difference statistic (i.e., the number of simulations in the magenta region, divided by the total number of simulations ($10^8$)).

**Movie S1. Representative time-lapse of *C. elegans* germline and embryo expressing *rps-27*>DHB::2x-mKate2,** Related to Figure 1, S1. Time-lapse (*n*=4 germline per genotype examined) following the division of germline nuclei expressing H2B (top) and DHB (bottom). The time points were acquired every 3 min for 78 min. The video was constructed from a single confocal section. Time-lapse (*n*=1 embryo per genotype examined) following cell divisions in the early *C. elegans* embryo expressing H2B (middle), DHB (right) and the overlay (left). The time points were acquired every 3 min for 45 min. Scale bar = 10 μm.

**Movie S2. Representative time-lapse of S to G2 transition during sex myoblast (SM) division expressing *rps-27*>DHB::2xmKate2 and *pcn-1*>PCN-1::GFP,** Related to Figure 1, S1. Time-lapse (*n*=7 animals examined). Compiled movie follows cell cycle progression visualizing both the CDK sensor (top) and PCNA (bottom). Loss of PCNA puncta in frame 16 correlates with G2 phase entry. The time points were acquired every 5 min for 125 min. The video was constructed from single confocal Z-sections (1 μm step size) selected as a sub-stack from 20 total Z positions. Scale bar = 10 μm.

**Movie S3. Representative time-lapse of pre-terminal and terminal sex myoblast (SM) division expressing *rps-27>DHB::GFP*,** Related to Figure 2, S2. Time-lapse (*n*=5 pre-terminal and *n*=3 terminal animals examined). Compiled movie follows the first two pre-terminal divisions of the SM cells (white arrowheads). The time points were acquired every 5 min for 300 min. The second movie follows the third and terminal division of the SM cells (white arrowheads). The time points were acquired every 5 min for 185 min. The video was constructed from single confocal Z-sections (1 μm step size) selected as a sub-stack from 20 total Z positions. Scale bar = 10 μm.

**Movie S4. Representative time-lapse of pre-terminal and terminal uterine sheath (SS) divisions expressing *rps-27>DHB::GFP*,** Related to Figure 2, S2. Time-lapse (*n*=7 pre-terminal and *n*=4 terminal animals examined). Compiled movie follows a pre-terminal division of the SS cells (white arrowheads). The time points were acquired every 5 min for 240 min. The second movie follows a terminal division of the sheath cells (white arrowheads). The time points were acquired every 5 min for 180 min. The video was constructed from single confocal Z-sections (1 μm step size) selected as a sub-stack from 20 total Z positions. Scale bar = 10 μm.

**Movie S5. Representative time-lapse of pre-terminal and terminal vulval precursor cell (VPC) divisions expressing *rps-27>DHB::GFP*,** Related to Figure 3, S2. Time-lapse (*n*=15 pre-terminal and *n*=10 terminal animals examined). Compiled movie follows a pre-terminal division of the VPC cells through the birth of the terminal D cells. The time points were acquired every 5 min for 240 min. The second movie follows the terminal divisions of the majority of the remaining VPCs. The time points were acquired every 5 min for 175 min. The video was constructed from single confocal Z-sections (1 μm step size) selected as a sub-stack from 20 total Z positions. Scale bar = 10 μm.

**Movie S6. Representative time-lapse of vulval D cell division expressing *rps-27>DHB::GFP***, Related to Figure 5, S4. Time-lapse (*n*=10 animals examined), following the division of C (blue *) and D (green *) VPC cells. The time points were acquired every 5 min for 270 min. The video was constructed from single confocal Z-section sections (1 μm step size) selected as a sub-stack from 20 total Z positions. Scale bar = 10 μm.

**Movie S7. Representative time-lapse of birth of CDK$^{low}$ cells in the posterior growth zone of 22 somite stage zebrafish,** Related to Figure 6. Time-lapse (*n*=1 animal, 16 cell births tracked). Time points were acquired every 5 min for three hours. The video was constructed from a single confocal Z-section. Yellow and blue arrows follow the birth of two cells that are born into a CDK$^{low}$ state (DHB cytoplasmic:nuclear ratio shown in first and last frames). Green shows membrane marker LCK.mNeonGreen (Lck) and magenta shows DHB.mScarlet (DHB). Scale bar = 10 μm.
